# Beyond efficacy: Persistence and off-target effects of three biological nitrification inhibitors in two contrasting agricultural soils

**DOI:** 10.64898/2026.07.11.737981

**Authors:** Paula A. Rojas-Pinzon, Bernhard Seidl, Stella Kejik, Christopher J. Sedlacek, Judith Prommer, Christoph Bueschl, Taru Sandén, Heide Spiegel, Andrew T. Giguere, Lucia Fuchslueger, Petra Pjevac

## Abstract

The use of nitrogen (N) fertilizers to meet global food demands is expected to continue rising. However, up to 70% of N applied to agricultural soils is lost through microbially mediated processes such as nitrification. Inhibiting nitrification is thus a key strategy to reduce N losses and improve fertilizer N use efficiency. Various plant-derived compounds, termed biological nitrification inhibitors (BNIs), have been shown to reduce accumulation of nitrification products, intermediates, and byproducts (nitrite, nitrate, nitric and nitrous oxides). However, the mechanisms by which BNIs affect nitrifiers, along with their specificity and persistence in soil are not well understood. Here, we evaluated the effects of three BNIs: methyl 3-(4-hydroxyphenyl) acrylate (MHPA), 6-methoxy-2(3H)-benzoxazolone (MBOA), and limonene, on ammonia-oxidizing, total microbial, and fungal communities in two soils with contrasting pH. Their persistence in each soil was also evaluated. Although ammonia-oxidizing archaea initially dominated nitrifier communities in both soils, their bacterial counterparts significantly increased after mineral N addition but also were more sensitive to BNI application. Limonene and the synthetic inhibitor DMPP stimulated ammonium immobilization, as total soil mineral N was significantly reduced. Limonene and MHPA had the strongest off-target effects, increasing the relative abundance of hydrocarbon-degrading bacteria and potential fungal pathogens, respectively. In contrast, MBOA inhibited nitrification with minimal off-target effects. Among the tested BNIs, MBOA was also the most persistent in the high-pH, high-nitrification-rate soil. Our results show that MBOA is a promising biological inhibitor and highlight the importance of understanding BNIs’ ecological effects to develop targeted and sustainable N management strategies.

## 1. Introduction

Recent projections estimate that by 2050 the global population will reach 10 billion and will require a ∼60% increase in food production (van Dijk et al., 2021; Alexandratos and Bruinsma, 2012). To meet these needs, the use of nitrogen (N) based fertilizers in agricultural systems is also projected to continue rising (Rohr et al., 2019). Currently, the fertilizer N use efficiency (NUE) of agricultural systems varies considerably, with losses reaching up to ∼70% of the total N applied (Tilman et al., 2002; Hegedus et al., 2023). In addition, agricultural systems contribute significantly to global greenhouse gas emissions, accounting for ∼80% of anthropogenic nitrous oxide (N2O) emissions (Birch, 2014). Low NUE and high N losses reduce crop yields, increase crop production costs and have significant environmental impacts (West et al., 2014; Gu et al., 2023; He et al., 2024). Therefore, improving NUE is essential to ensure that future food demands are met with a lower environmental impact.

Whether applied as NH_3_/NH_4_^+^ or converted to NH_3_/NH_4_^+^ through urea hydrolysis, fertilizer N is highly susceptible to oxidation by nitrifying microorganisms (Subbarao et al., 2013). Nitrification is the first step in the cascade of N transformations leading to N losses from agricultural soils. Four groups of autotrophic microorganisms are involved in the sequential oxidation of ammonia (NH_3_) to nitrate (NO_3_^-^): ammonia-oxidizing archaea (AOA) and bacteria (AOB), which oxidize NH_3_ to nitrite (NO_2_^-^), nitrite-oxidizing bacteria (NOB) which oxidize NO_2_^-^ to NO_3_^-^ and, complete ammonia-oxidizing (comammox) bacteria, which perform the complete oxidation of NH_3_ to NO_3_^-^ (Kuypers et al., 2018). While NH₄⁺ is predominantly soil-bound, NO₂⁻ and NO₃⁻ (further on referred to together as NOₓ⁻), are highly mobile and easily leached into groundwater, contributing to eutrophication (Norton and Ouyang, 2019). Additionally, NH₃ oxidation and NOₓ⁻ reduction during denitrification contribute to the formation of the potent greenhouse gas nitrous oxide (N₂O), whose release into the atmosphere amplifies the overall environmental impact of agricultural N losses.

Numerous microcosm and field studies have demonstrated that reducing soil nitrification rates is a promising management strategy to reduce NO_3_^-^ leaching (Di and Cameron, 2004; Shi et al., 2016), N_2_O emissions (Yin et al., 2023), and increase plant N uptake (Pasda et al., 2001; Sun et al., 2016; Karwat et al., 2017). Several synthetic nitrification inhibitors (SNIs) are commercially available and are applied together with N-based fertilizers (Subbarao et al., 2006; Zerulla et al., 2001). In Europe, the most commonly used SNI is 3,4–dimethylpyrazole phosphate (DMPP) (Biewald et al., 2025), which can be applied at lower concentrations (1% of the applied N) than other commercially available SNIs, such as dicyandiamide (DCD) or nitrapyrin. Similar to other SNIs, the efficacy of DMPP is influenced by many environmental factors (Guardia et al., 2018). It is highly soluble and prone to leaching (Marsden et al., 2016), and conflicting results regarding off-target effects have been reported (Kong et al., 2016; Rime and Niklaus, 2017; Bachtsevani et al., 2021). Consequently, current research efforts focus on discovering novel inhibitors, including naturally occurring, plant-derived alternatives, known as biological nitrification inhibitors (BNIs).

A large diversity of BNI compounds have been identified in tissue and root exudates of a variety of plants (Subbarao et al., 2012, 2013, 2015; Coskun et al., 2017). Several have been shown to inhibit soil nitrification including limonene (White, 1991), methyl 3-(4-hydroxyphenyl) acrylate (MHPA) also known as methyl-*p*-coumarate (Gopalakrishnan et al., 2007), and 6-methoxy-2(3H)-benzoxazolone (MBOA) (Otaka et al., 2021, 2023). Previous studies with the AOB *Nitrosomonas europaea*, showed reduced nitrite production in the presence of these BNIs (Ward et al., 1997; Gopalakrishnan et al., 2007; Otaka et al., 2023). Other studies have also used the decrease in NO_3_^-^ and/or NO_2_^-^ accumulation as a proxy for the efficacy of known and newly identified BNIs (Subbarao et al., 2013; Nuñez et al., 2018; Otaka et al., 2023). However, compared to SNIs, little is known about the effects of BNIs on the activity of different ammonia-oxidizing guilds, their modes of action, their target(s) of inhibition, their off-target effects, and their persistence in agricultural soils (Gopalakrishnan et al., 2009; Lu et al., 2019; Ma et al., 2021; Kaur-Bhambra et al., 2022; Lan et al., 2022; Kolovou et al., 2023).

The mode of action and inhibition target assessment of inhibitors are usually performed using model organisms in pure culture experiments (Papadopoulou et al., 2020; Kaur-Bhambra et al., 2022; Yildirim et al., 2024). However, off-target effects, such as an increase in soil respiration (Rojas-Pinzon et al., 2024), nitrous oxide reductase (*nosZ*) gene abundance and transcription (Torralbo et al., 2017; Friedl et al., 2020), and inhibition of gross N mineralization (Lan et al., 2023) have been observed after NI addition to soils. The persistence of BNIs in soil determines not only their efficacy, but also the extent to which they can accumulate in soil, which could lead to unwanted long-term effects. Soil properties, as well as the chemical properties of the NIs, play an important role in NI persistence (Nardi et al., 2013; Marsden et al., 2016; Guardia et al., 2018). For instance, some SNIs can accumulate in higher trophic levels due to their high mobility and resistance to degradation (Macadam et al., 2003; Chen et al., 2014; Pal et al., 2016). In addition, a decrease in mobility and efficacy has been observed in NIs applied to soils with high organic matter and clay content (Puttanna et al., 1999; McGeough et al., 2016). While several studies have examined the effects of BNIs on the soil nitrifying community (Nardi et al., 2013; Lu et al., 2019), comparatively few have investigated the possible effects of BNI addition on other soil carbon and nitrogen transformation processes (Florio et al., 2021; Wang et al., 2021) or determined the fate and persistence of the BNIs themselves (Ma et al., 2021).

The objective of this study was to further evaluate the specificity and persistence of three BNIs (limonene, MHPA, and MBOA) in agricultural soils, compared to the widely used SNI, DMPP. Soil microcosms with two agricultural soils of contrasting pH were employed to assess the effect of NIs on the nitrifying microbial communities, off -target effects on bacterial and fungal communities, as well as NI persistence over a period of several weeks. While previous studies have focused on the effect of BNIs on NO_x_^-^ or N_2_O production across different soil types, this study provides additional insights into BNI effects beyond their capacity to suppress soil nitrification, which is crucial for evaluating their overall suitability for field application.

## 2. Materials and methods

### 2.1. Study sites

Two agricultural soils from long-term fertilization experiments managed by the Austrian Agency for Health and Food Safety (AGES) were sampled in February 2024. The sites are located in the Marchfeld region of Lower Austria (48°12’57.2’’N 16°37’06.1’’E) and in the Alpenvorland region (48°07’31.7’’N 15°09’13.4’’E). The Marchfeld soil is classified as Calcaric Phaeozem (sandy loam: 30.3% sand, 45.6% silt, and 24.2% clay) with a pH in water of 8.50. This alkaline soil (AS) had an initial NH_4_^+^ content of 0.1 ± 0.02 µg N g^-1^ soil and a NO_3_^-^ content of 7.7 ± 0.09 µg N g^-1^ soil and has a cation exchange capacity of 27 ± 0.41 cmolc kg^-1^. The soil from the Alpenvorland site is a Gleyic Luvisol (loamy silt: 9.5% sand, 71.2% silt, and 19.4% clay), has a circumneutral soil (CS) pH in water of 6.12 and a cation exchange capacity of 8.8 ± 0.36 cmolc kg^-1^. The initial NH_4_^+^ content was 0.2 ± 0.06 µg N g^-1^ soil, and the NO_3_^-^ content was 3.0 ± 0.06 µg N g^-1^ soil.

The long-term N and phosphorus fertilization for both soils have consisted of 120 kg N ha^−1^ yr ^−1^ and 75 kg P_2_O_5_ ha^−1^ yr ^−1^, respectively. The crop rotations consisted of ∼55% cereals and ∼45% root crops (Lehtinen et al., 2014; Spiegel et al., 2018). Both sites were uncropped at the time of sampling. A single composite soil sample was collected from each site by combining ten individual soil cores from the top 10 cm of soil. The soil samples were transported to the laboratory, sieved (2 mm mesh size), and stored at field temperature (10°C). Soil for molecular analysis was stored at -80°C.

### 2.2. Microcosm incubations

Soil microcosms containing 2 g of soil were set up in 27 ml serum bottles. The soil water content was adjusted to 60% water holding capacity (WHC) by evenly distributing a solution of ammonium sulfate to achieve a final concentration of 130 μg (NH_4_)_2_SO_4_–N g^-1^ dry weight (dw) soil. The treatments consisted of a control that did not receive any nitrification inhibitor, and microcosms that received DMPP, MHPA, MBOA, or limonene. DMPP was applied at a rate of 1% of the applied N (1.3 μg g^-1^ dw soil) as part of the (NH_4_)_2_SO_4_ addition. MBOA, MHPA, and limonene were premixed with the soil prior to (NH_4_)_2_SO_4_ addition due to their low water solubility. A set of different MHPA and limonene concentrations were previously tested in nitrification potential assays in soils from these sites (Rojas-Pinzon et al., 2024). The same nitrification potential experiments as performed in Rojas-Pinzon et al. (2024), were here repeated to determine the inhibitory potential of MBOA (Fig. S1). The concentrations resulting in 80% nitrification inhibition were applied in the microcosms. MBOA was applied at a rate of 290 μg g^-1^ dw soil for the AS and 111 μg g^-1^ dw soil for the CS. MHPA was applied at rates of 387 and 394 μg g^-1^ dw soil for the AS and the CS, respectively. Limonene was applied at rates of 1.8 mg g^-1^ dw soil for the AS and of 786 μg g^-1^ dw soil for the CS. The serum bottles were sealed with butyl rubber stoppers and aluminum caps and incubated in the dark at 20°C. For each soil a set of 20 bottles were prepared and destructively sampled across five time points (four replicates per time point).

Previous incubations showed higher nitrification activity in the AS than in the CS (Rojas-Pinzon et al., 2024). To account for this, incubations with the AS were sampled on days 0, 2, 3, 8 and 15, while the CS was sampled on days 0, 3, 6, 15 and 21. At each time point a subsample of soil (0.5 g) was stored at -80°C for molecular analyses. The remaining sample (∼ 1.5 g) was mixed with 15 mL 2 M KCl for mineral N extraction. NH_4_^+^ consumption over time was quantified by a modified photometric indophenol reaction method (Mulvaney, 1996), while the acidic VCl_3_/Griess reaction was used to determine NO_2_^−^ and NO_3_^−^ accumulation (Miranda et al., 2001).

### 2.3. Quantification of *amoA* gene, transcripts and amplicon sequencing

Total nucleic acids were extracted from 0.5 g of soil using a modified phenol-chloroform method (Angel et al., 2012) and purified using the OneStep PCR inhibitor Removal kit (Zymo Research, USA). An aliquot was stored for DNA-based gene amplification, sequencing and qPCR, while the remaining extract was treated with the Turbo DNA-free kit (Thermo Fisher Scientific, USA) to deplete DNA. The purified RNA served as template for RT-qPCR, performed using the Luna Universal One-Step RT-qPCR kit (New England Biolabs, USA). Gene and transcript copy numbers of the ammonia monooxygenase subunit A (*amoA*) gene of AOB, AOA, and comammox were quantified for samples collected at days 0, 3, 8 and 15 for the AS and at days 0, 6, 15 and 21 for the CS. Reagents, primers, as well as qPCR and RT-qPCR conditions are listed in Table S1. A control without reverse transcriptase was included in all cases to exclude residual DNA contamination. Standard curves for each gene were generated from serial dilutions of 10^7^–10^1^ gene copies µl^−1^ of linearized plasmids with insertions of the target genes. All transcript quantifications were performed in duplicate in a final reaction volume of 20 µl.

Amplification and sequencing of the bacterial and archaeal 16S rRNA gene, fungal ITS2 region, and clade-specific *amoA* genes were performed on samples collected at day 0 and at the end of the experiment (day 15 for the AS and 21 for the CS), following a two-step PCR protocol (Pjevac et al., 2021) using the primers listed in Table S1.

Sequencing was performed on the Illumina MiSeq platform at the Joint Microbiome Facility of the Medical University of Vienna and the University of Vienna JMF (project ID: JMF-2411-07), using the 600-cycle v3 chemistry (2 × 300 bp paired-end reads). Amplicon sequence variants (ASV) were inferred using the DADA2 pipeline (Callahan et al., 2016) following the recommended workflow. The taxonomy of the prokaryotic 16S rRNA gene amplicons was assigned using the SILVA database SSU Ref NR 99 release 138.1 (Quast et al., 2013), and the UNITE database (Abarenkov et al., 2020) was used for classification of fungal ITS2. For AOA, AOB, and comammox clade B *amoA* taxonomy assignment, custom databases were created by retrieving the respective *amoA* entries from NCBI GenBank.

### 2.4. Nitrification inhibitor recovery

To evaluate the persistence of the NIs in soil, another set of microcosm incubations was established exactly as described above. Soil microcosms were destructively sampled at days 0, 3, 8, and 15 for the AS and 0, 6, 15, and 21 for the CS. Samples were then frozen at -80°C until NI extraction. DMPP, MHPA and MBOA extractions were performed following protocols developed by Adhikari et al. (2021). In brief, for DMPP extraction, 2 mL of HPLC eluent (212 mL methanol mixed with 788 mL of a 1 mM H_3_PO_4_ and 5 mM KH_2_PO_4_ buffer) and 8 mL of 2 M KCl were added to 2 g of fresh soil. This solution was shaken (1 h at 200 rpm) and centrifuged (5 mins at 13,751 *g*). The resulting supernatant (10 mL) was mixed with 400 μL 1 M NaOH and 8 mL di-ethylether. After a second round of shaking (10 min at 200 rpm) and centrifugation (10 min at 13,751 *g*), 5 mL of the supernatant was transferred to a new tube containing 2 mL of HPLC eluent. Diethyl ether was allowed to evaporate overnight, and the samples were stored at -80°C until analysis. For MHPA and MBOA the extraction protocol for non-water-soluble NIs by Adhikari et al. (2021) was followed with some modifications. Soil (2 g) was first mixed with 4 mL of HPLC-grade acetonitrile, shaken on an orbital shaker (30 min at 220 rpm). The mixture was centrifuged (10 min at 13,751 *g*), and 4 mL of the resulting supernatant was filtered through a 0.45 μm filter and stored at -80°C. Limonene was extracted using the same protocol, but with n-hexane instead of acetonitrile.

Quantification of BNIs in acetonitrile extracts and of DMPP in the HPLC eluent was performed with an IQ-X Orbitrap equipped with a heated electro-spray ionization (H-ESI) source coupled to a Vanquish Horizon UHPLC system (Thermo Fisher Scientific, San Jose, CA, USA). Limonene in n-hexane extracts was analyzed following the procedure established by Huang et al. (2022) using gas chromatography (7890 B Agilent Technologies, USA) connected to a time of flight (TOF) MS (Pegasus BT, LECO, USA) equipped with a split/splitless injector and a DB-5 column (60 m; Agilent, USA). Between five and eight concentrations of each compound dissolved in the respective extraction solvent were used to generate standard calibration curves. The Thermo Fisher Scientific Software Xcalibur QuanBrowser (Version 4.7.69.37), the Avalon peak-picking algorithm and the ChromaTOF software (Version 5.56.53) were used for data evaluation. Operating conditions for the Orbitrap and gas chromatography measurements, as well as the calculations used to determine the amount of NI recovered in each soil over time, are specified in the supplementary material.

### 2.5. Statistical analyses

All statistical analyses were performed in R versions 4.4.2 and 4.4.3 (R Core Team, 2024) using RStudio server version 2024.12.1. To evaluate the effects of NI addition and incubation time on NH₄⁺ consumption, NO₃⁻/NO₂⁻ accumulation, and *amoA* transcription, a two-way ANOVA followed by a post-hoc Tukey test was used when data were normally distributed and variances were homogeneous (detailed statistical tests are shown in the supplementary material). When data did not meet these assumptions, a generalized linear model (*glm* function) was used, followed by pairwise comparisons of estimated marginal means (EMMs) with Bonferroni correction via the *emmeans* function of the *emmeans* 1.11.1 package (Lenth, 2025). Differences in NI concentration and in the percentage of NI recovered over time were assessed with a one-way ANOVA followed by a Tukey test for normally distributed data with homogeneous variances, or with a Kruskal–Wallis test followed by Dunn’s test when assumptions were not met.

Amplicon sequencing data were analyzed using the *phyloseq* 1.50.0 (McMurdie and Holmes, 2013), *microbiome* 1.28.0 (Lahti and Shetty, 2012), and *DEseq2* 1.46.0 (Love et al., 2014) packages. Alpha-diversity analysis was performed on rarefied datasets. To assess the effect of the NIs on microbial community composition, principal component analysis (PCA) on an Aitchison distance matrix was performed separately for each amplicon dataset from each soil. Finally, potential off-target effects of NI application on 16S rRNA- and ITS2-based microbial communities were assessed through genus-level differential relative abundance analyses, after excluding genera with an average relative abundance < 2% in any treatment.

## 3. Results

### 3.1. Mineral nitrogen dynamics in response to NI amendments

To evaluate the impact of BNI amendments on nitrification in fertilized agricultural soils, the dynamics of mineral N were monitored using soil microcosms. In control microcosms with no NI addition, a faster ammonia oxidation rate was observed in the AS than in the CS. In the AS, all NH_4_^+^ was oxidized to NO_2_^-^ and NO_3_^-^ (NO_x_^-^) within 15 days, while in the CS, approximately 27 ± 2.6 µg g^-1^ dw soil of NH_4_^+^ remained after 21 days (Fig. 1A, Table S2). In both soils, the addition of the SNI DMPP resulted in the highest amount of soil-retained NH_4_^+^ at the end of the incubation period, followed by the BNI MBOA (Fig. 1A, Table S2). Limonene only increased soil NH_4_^+^ retention significantly on day 15 in the AS, and was not an effective inhibitor in the CS. In contrast, MHPA was not an effective inhibitor, irrespective of soil type (Fig. 1A).

**Fig. 1.**
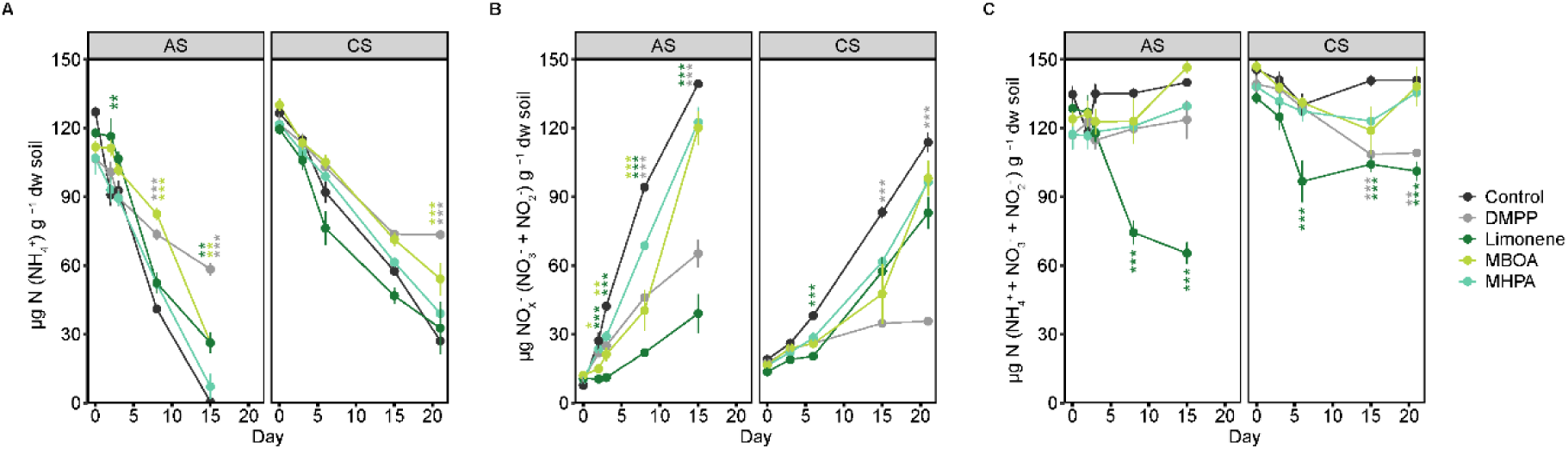
Mineral N consumption and production in the AS and CS with and without addition of NIs. (A) NH_4_^+^ consumption, (B) NO_X_^-^ (NO_3_^-^ and NO_2_^-^) accumulation, and (C) total inorganic N (NH_4_^+^, NO_3_^-^ and NO_2_^-^) recovery over 15 days for the AS and 21 days for the CS. Data are normalized per gram dry weight (dw) soil and presented as means ± SE (n = 4). Asterisks and the color denote significant differences of a specific treatment relative to the control on each day (* < 0.05, ** < 0.01, *** < 0.001). Post -hoc test results of differences between treatments on each day for each soil are provided in Tables S4-S6.

In addition to soil NH_4_^+^ dynamics, the inhibition of NO_x_^-^ accumulation is also a determinant of nitrification inhibition (Fig. 1B). DMPP significantly inhibited NOx-accumulation compared to control incubations in both soils (Fig. 1B) and was the only inhibitor to significantly decrease NO_x_^-^ accumulation in the CS by the end of the 21-day incubation period. Both MBOA and limonene significantly inhibited NO_x_^-^ accumulation in the AS. MBOA showed an inhibitory effect early in the incubation with its efficacy diminished by day 8, whereas limonene had the lowest NO_x_^-^ accumulation compared to the control from day 2 until the end of the 15-day incubation period (Fig. 1B). Similar as for the NH_4_^+^ content, MHPA was the least effective inhibitor of NO_x_^-^ accumulation in the AS. Although limonene significantly inhibited NOx⁻ accumulation after 6 days of incubation, by the end of the incubation period, none of the three BNIs significantly inhibited NOx⁻ accumulation in the CS microcosms.

In the absence of NIs, the average amount of total inorganic N recovered across all timepoints (NH4^+^, NO3^-,^ and NO2^-^) was 100 ± 1.1 % in the AS, and 100 ± 1.2 % in the CS (Fig. 1C). These stable recoveries of total inorganic N over time suggest little influence of microbial N transformations other than nitrification in the microcosms. A balanced inorganic N recovery was also observed in microcosms with MBOA (97. 1 ± 2.4 % in the AS and 96.4 ± 2.2 % in the CS), MHPA (91.1 ± 1.7 % in the AS and 93.9 ± 1.1 % in the CS), and DMPP in the AS (90.5 ± 2.1 %). However, the microcosms with DMPP in the CS and limonene in both soils resulted in less total inorganic N recovered compared to the control (89.3 ± 2.4 % with DMPP in the CS, 78.4 ± 5.6 % for limonene in the AS and 80.0 ± 2.5 % in the CS. Fig. 1C). This imbalance between the NH_4_^+^ consumed and the corresponding NO_x_^-^ accumulated occurred after six days with DMPP in the CS, and after three days with limonene in both soils.

### 3.2. Abundance and transcription of *amoA* genes

To evaluate the effects of NIs on ammonia-oxidizing communities, AOB, AOA, and commamox *amoA* gene and transcript copy numbers were quantified. In both soils, the native nitrifier communities were AOA dominated, with the average AOA *amoA* gene copy numbers being an order of magnitude higher (5.6 ± 0.5 x 10^6^ copies g^-1^ dw soil in the AS, and 2.9 ± 0.2 x 10^6^ copies g^-1^ dw soil in the CS) than AOB (5.1 ± 0.3 x 10^5^ copies g^-1^ dw soil in the AS, and 2.3 ± 0.1 x 10^5^ copies g^-1^ dw soil in the CS) or the average comammox clade B *amoA* gene copy numbers (9.2 ± 0.5 x 10^4^ copies g^-1^ dw soil in the AS, and 3.7 ± 0.1 x 10^5^ copies g^-1^ dw soil in the CS) (Table S3, Fig. S2). In both soils and across all treatments, gene abundances significantly decreased or remained unchanged over the incubation period for AOA *amoA* (0.3-0.9-fold change). In contrast, AOB *amoA* gene abundance changes were NI treatment-specific: AOB *amoA* gene abundance significantly increased in the control, MBOA, and MHPA treatments (3.3-5.6-fold change), but did not change significantly in the limonene treatment in both soils or in the DMPP treatment in the AS (1.2-2.2-fold change) (Table S3, Fig. S2). Overall, comammox clade B *amoA* abundances were stable throughout the incubation period, with small significant changes that were specific to each soil and NI treatment (0.6-2.6-fold change) (Table S3, Fig. S2).

Initially, AOA *amoA* transcripts had the highest abundance in both soils (2.38 ± 0.1 x 10^5^ and 3.89 ± 0.1 x 10^5^ transcript copies g^-1^ dw soil in the AS and the CS, respectively) (Fig. 2A). The abundance of AOB *amoA* transcripts was higher in the AS than in the CS (1.49 ± 0.1 x 10^4^ and 1.33 ± 0.1 x 10^3^ transcript copies g^-1^ dw soil, respectively) (Fig. 2B). Although comammox clade B *amoA* gene copies were detected in both soils, no comammox clade B *amoA* transcripts were detected in either soil across any treatment. The *amoA* transcription patterns of AOA and AOB differed between soils but followed similar overall trends in response to the NI treatments.

**Fig. 2.**
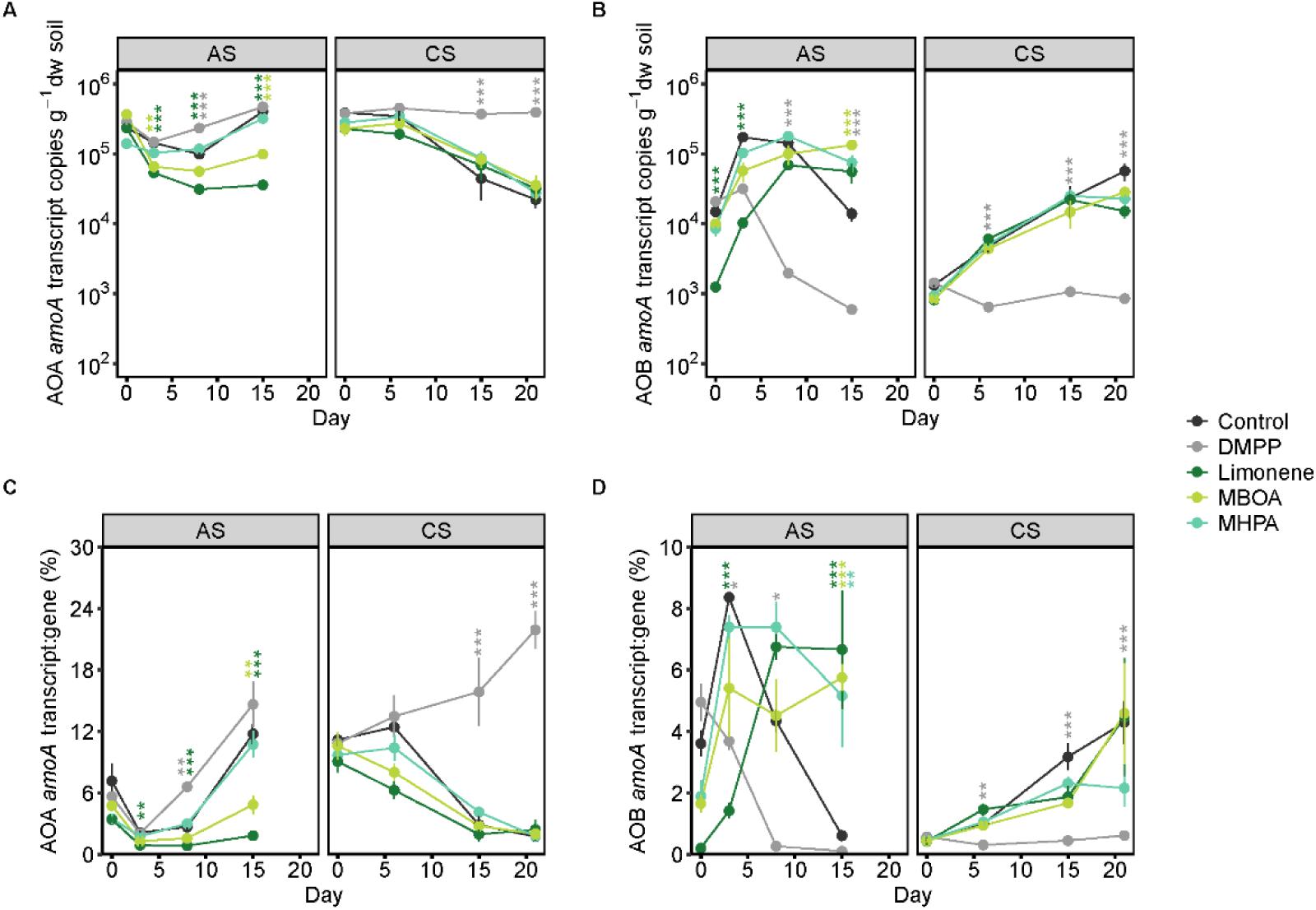
Change in transcript abundance and transcript-to-gene ratio of the AOA *amoA* (A and C) and the AOB *amoA* genes (B and D) in the AS and CS with and without addition of NIs. Data are normalized per gram dry weight (dw) soil and presented as means ± SE (n = 4). Asterisks and the color denote significant differences of a specific treatment relative to the control on each day (* < 0.05, ** < 0.01, *** < 0.001). Post-hoc test results of differences between treatments on each day for each soil, are provided in Tables S7-S10. Gene abundances are provided in Table S3 and Fig. S2.

In the AS, the NH_4_^+^ addition, both with and without NIs, was followed by an immediate decrease in AOA *amoA* transcript abundances, before stabilizing at a significantly lower level (limonene and MBOA), or subsequently increasing to pre-treatment abundances at later timepoints (control, DMPP, and MHPA) (Fig. 2A). The AOB *amoA* transcript abundances showed an inverse trend where an initial increase followed by a continuous recovery to pre-treatment abundances was observed (control, MHPA and limonene) (Fig. 2B). The MBOA treatment showed a similar transcript abundance trajectory but stabilized at a significantly higher AOB *amoA* transcript abundance at day 15, compared to control microcosms. In contrast, DMPP suppressed the increase in AOB *amoA* transcript abundances after NH_4_^+^ addition, and significantly reduced AOB *amoA* transcript abundances compared to the control after day 8 (Fig. 2B). In the CS, AOA *amoA* transcript abundances decreased in the control and all three BNIs treatments (Fig. 2A). In contrast, AOB *amoA* transcript abundances increased throughout the incubation period in the control and all three BNI treatments. In the DMPP treated CS microcosms, AOA and AOB *amoA* transcript abundances remained stable over the entire course of the incubation and were significantly different than all other treatments (Fig. 2A, B).

Not only did AOA outnumber AOB in both soils (Table S3, Fig. S2), but they also had a higher transcript to gene ratio in both soils before treatment (Fig. 2C, 2D). The transcript to gene ratios for the AOB and AOA also showed an inverse pattern across all treatments, similar to the transcript abundances. In the AS, where the AOA transcript to gene ratio in all treatments initially decreased between days 0-3 (0.21 ± 0.03 - 0.54 ± 0.10-fold-change) before increasing until day 15 (1.22 ± 0.16 - 5.02 ± 1.3-fold-change. Fig. 2C). In contrast, the transcript to gene ratio for AOB across the control and three BNI treatments increased between days 0-3 (2.43 ± 0.33 - 7.16 ± 1.52-fold-change), before stabilizing or decreasing until day 15. In the CS, this general trend is observed as a steady decrease in the AOA transcript to gene ratio (in the control and three BNI treatments), while the opposite occurred for the AOB transcript to gene ratio (Fig. 2C, D). In contrast, the addition of DMPP in the CS resulted in a significant increase in the AOA transcript to gene ratio and a consistently low AOB transcript to gene ratio throughout the incubation period.

### 3.3. Effect of the NIs on ammonia oxidizer communities

While similar richness and evenness was observed in the AOA and AOB communities of the AS and CS by *amoA* gene amplicon sequencing, the commamox clade B communities showed lower diversity indices (Fig. S3). Overall, the richness and evenness of the ammonia oxidizer communities in the AS remained unaffected regardless of treatment. In the CS, community richness and evenness changes were NI treatment specific. In the control and MBOA microcosms significantly higher AOA but lower AOB and comammox clade B richness and evenness was observed after the incubation (Fig. S3). Limonene had the same effect on AOA and comammox clade B communities, while the AOB community remained unaffected (Fig. S3). Interestingly, DMPP treatments had no effect on the richness and evenness of any of the ammonia oxidizer communities (Fig. S3). Regardless of the type of NI applied, the AOA communities in both soils showed no pronounced compositional shifts (Fig. S4). In contrast, MHPA, MBOA, and limonene applications resulted in AOB community divergence from the initial and control microcosm compositions (Fig. S4). Of the three BNIs tested, only limonene caused a compositional shift in the commamox clade B communities in both the AS and CS (Fig. S4).

### 3.4. Off-target effects

Differential relative abundance analyses of the total microbial communities, based on 16S rRNA gene amplicon sequencing, showed that BNI application led to a significant increase in the relative abundance of several bacterial genera, but to no taxon-specific suppression in either soil (Fig 3A). More bacterial genera were responsive to BNI application in the AS compared to the CS. Limonene and MHPA caused a pronounced effect in both soils, leading to an increase in the relative abundance of genera including known hydrocarbon degraders and denitrifiers (*Rhodococcus* and *Pseudomonas*), methylotrophs (*Methylovorus* and *Methylotenera*), as well as chitinolytic bacteria (*Chitinophaga*). MBOA application exhibited a minor effect by only increasing the abundance of one genus of methylotrophs in the AS (*Methylovorus*). Notably, limonene addition led to an extreme proliferation of *Rhodococcus*-related microorganisms in both soils, with their relative abundance reaching 30% and 50% in the CS and AS, respectively. This *Rhodococcus* bloom resulted in a significant reduction in the overall soil richness and evenness, as well as in greater dissimilarities in the total microbial community composition after incubation of soils with limonene. In contrast, the addition of MHPA, MBOA and DMPP did not significantly alter the total microbial community alpha and beta diversity (Fig. S5 and S6).

**Fig. 3.**
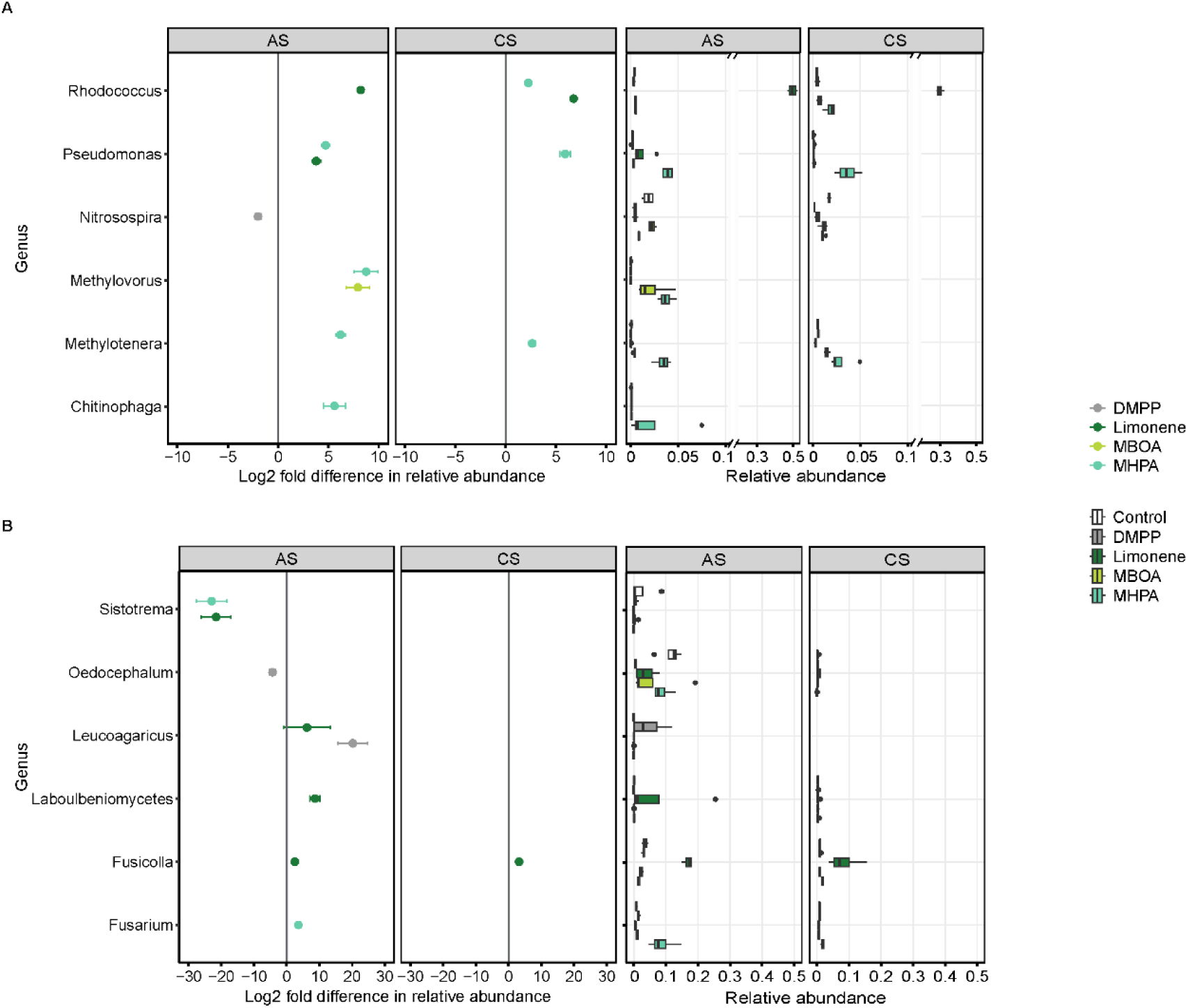
Genus-level differential relative abundance changes for the (A) 16S rRNA and (B) ITS gene-based microbial community compositions after 15 and 21 days incubation with and without NIs for the AS and the CS, respectively. Only genera with a relative abundance ≥ 0.02 in any of the treatments is depicted. Genera without relative abundance information in the CS is due to their absence in that soil. The median, first and third quartile of the relative abundances are depicted as the middle, upper, and lower hinges, respectively in the boxplots. The length of the whiskers is determined by the largest and the smallest value in the dataset that are within 1.5 times the interquartile range and outliers are shown as points (n = 4, p < 0.05). Data are presented as means ± SE (n = 4).

In both soils, fungal community composition was strongly affected by the BNI amendment compared to the 16S rRNA based microbial communities (Fig. 3B). Although the amendments had no significant effect on alpha diversity, BNI application resulted in greater community dissimilarity, particularly in the AS (Fig. 3B, S5 and S6). Similarly to the 16S rRNA gene-based microbial community, more fungal genera were responsive to BNI application in the AS compared to the CS, with limonene causing the greatest effects (Fig 3B). In both soils, mixed effects on the relative abundance of saprotrophic fungi and decomposers of complex organic compounds were observed, with increases or decreases depending on the specific NI applied. Notably, MHPA amendment resulted in a significantly increased relative abundance of the genus *Fusarium*, containing potential plant pathogens (Fig. 3B).

### 3.5. Recovery and persistence of NIs in soil

Sorption to soil particles and susceptibility to biotic and abiotic degradation have been shown to affect NI efficacy in soils (Marsden et al., 2016; McGeough et al., 2016; Guardia et al., 2018)). Therefore, the percent recovery of NIs as well as the change in recoverable NI concentration over time (i.e., NI persistence) in the AS and the CS were determined. Notably, from the start of the incubation only a fraction of the NIs added could be recovered from both soils. Limonene had the lowest initial percent recovery in both soils (16.0 ± 1.4 % in the AS and 3.4 ± 0.3 % in the CS, Table S11) despite being the NI applied at the highest concentration. Notably, MBOA had the highest initial percent recovery among all NIs in both soils (55.7 ± 4.4 % in the AS and 45.1 ± 4.2 % in the CS, Table S11). A significant reduction in recoverable NI concentrations was observed in both soils over time for all NIs (Fig. 4, Fig. S7). Amongst the BNIs, MBOA was the most persistent in the AS (36.3 ± 7.4% after 15 days), while all three BNIs persisted only until day 6 in the CS (Fig. 4). DMPP showed the highest persistence amongst all NIs with 53.9 ± 4.3 % in the AS and 43.2 ± 3.5 % in the CS at the end of the incubation period.

**Fig. 4.**
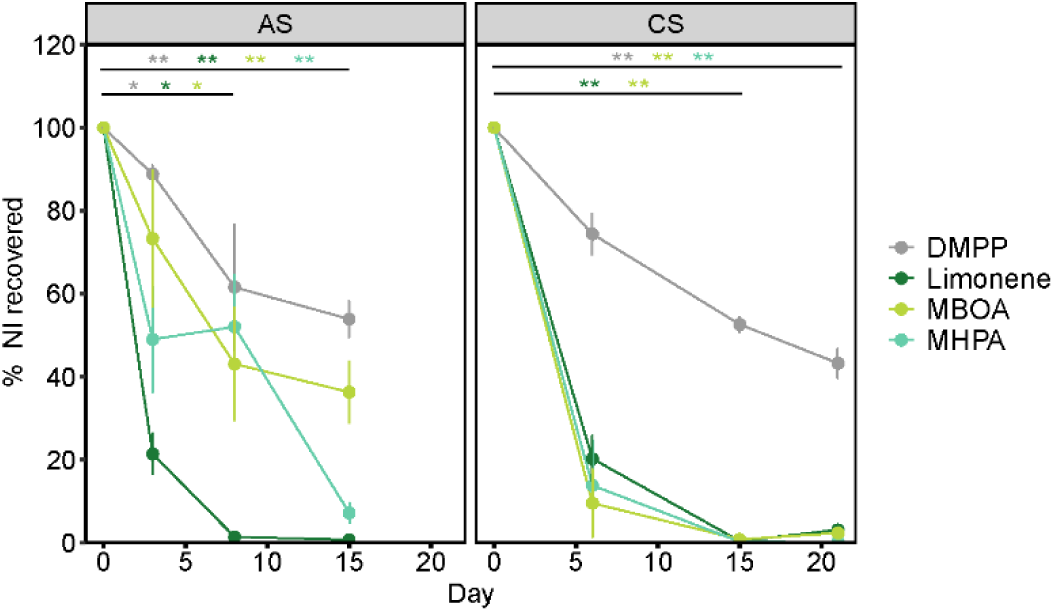
Persistence of NIs in the AS and CS. The percentage of NI recovered on each day and for each soil is depicted. Data are presented as means ± SE (n = 4). ANOVA or Kruskal-wallis test were used to assess statistical differences in the recovery of each NI over time. Asterisks and the color denote significant differences in the persistence of each NI over time (* < 0.05, ** < 0.01, *** < 0.001).

## 4. Discussion

The efficacy of NIs is often assessed by their ability to reduce NOx^-^ accumulation, which is typically associated with increased NH₄⁺ retention in soil and specificity toward inhibiting ammonia oxidizers (Subbarao et al., 2013; Nuñez et al., 2018; Otaka et al., 2023). However, whether NI application alters residual soil NH₄⁺, the extent to which NIs potentially affect non-nitrifying bacterial or fungal soil communities, and how these off-target effects are influenced by NI persistence in soil, remains unclear. In this study, we addressed these questions by examining the efficacy, specificity and persistence of three BNIs (limonene, MHPA, and MBOA) and the SNI DMPP in two contrasting agricultural soils.

### 4.1. Effects of NIs on total mineral N (NO_2_^-^ + NO_3_^-^ + NH_4_^+^)

Differences in soil physicochemical properties and in microbial community composition between the AS and the CS resulted in different nitrification rates. All of these factors have been shown to influence NI efficacy (Gilmour, 1984; Puttanna et al., 1999; Lu et al., 2019; Kuppe and Postma, 2024). A higher net nitrification rate was observed in the AS (8.7 ± 0.09 µg NO_X_^-^ g^-1^ dw soil day ^-1^) compared to the CS (4.5 ± 0.2 µg NO_X_^-^ g^-1^ dw soil day ^-1^), and yet limonene, MBOA and DMPP were more effective at reducing NO_x_^-^ accumulation in the AS (Fig. 1B). This is in contrast to a previous evaluation of these NIs in the same soils using potential nitrification assays instead of soil microcosms. Comparable net nitrification rates were observed with both methods; however, a higher inhibition efficacy was observed in the lower-pH CS soil (Rojas-Pinzon et al., 2024) (Figure S1). This discrepancy in results highlights the need for standardizing NI efficacy evaluation practices, and further acknowledgment of the known shortcoming of widely used nitrification potential assays (Taylor et al., 2010; Bachtsevani et al., 2021; Hazard et al., 2021; Rojas-Pinzon et al., 2024).

The SNI DMPP most efficiently inhibited nitrification in both soils, leading to significantly lower NO_x_^-^ accumulation and higher soil NH_4_^+^ retention (Figure 1A and B), supporting its established effectiveness as an SNI (Zerulla et al., 2001; Yang et al., 2013; Zhou et al., 2020). In contrast, MHPA — a hydrophobic compound — did not significantly inhibit nitrification in soil microcosms, regardless of soil type (Fig. 1A and B). A suboptimal concentration for soil microcosms, along with the influence of soil physicochemical properties, may have reduced MHPA’s efficacy compared to values determined in soil slurries. Although none of the three BNIs was more effective than DMPP, the results suggest that under certain soil conditions, BNIs such as MBOA may be suitable alternatives to SNIs.

NI application in agricultural soils is expected not only to reduce NO_x_^-^ accumulation, but also to increase and extend NH₄⁺ availability for crop uptake. Limonene in both soils and DMPP in the CS inhibited NO_X_^-^ accumulation (Fig. 1B), but this did not result in a corresponding elevated NH_4_^+^ content over time (Fig. 1A and C). Similar effects have been observed in microcosms treated with high concentrations of other BNIs, such as 1,9-decanediol and various terpenoids, and have been hypothesized to result from microbial nitrogen immobilization (Bremner and McCarty, 1988; Lu et al., 2019). Microbial N immobilization, potentially induced by a change in the C:N ratio of easily available substrates shifted by the applied NIs, or by a change in microbial nitrogen demand, might also have caused the N imbalance observed after limonene and DMPP application. Alternatively, this could be the result of only apparent inhibition, where the reduction in NOx-formation is due to enhanced microbial NO₃⁻ uptake rather than inhibition of nitrifiers. This has been previously observed and proposed as a potential apparent mode of action for some BNIs (White, 1991; Ma et al., 2021; Egenolf et al., 2022). Taken together, this highlights the importance of assessing NIs not only by their efficacy in reducing NOx⁻ accumulation, but also by their effects on total soil N dynamics.

### 4.2. Effect of NIs on ammonia oxidizers community and *amoA* expression

Contrasting results regarding the susceptibility of the different ammonia oxidizers to different NIs have been reported (Kaur-Bhambra et al., 2022; Kolovou et al., 2023; Yin et al., 2023; Issifu et al., 2024), highlighting the role of the physiological differences, and consequently of the soil ammonia oxidizer composition on the efficacy and specificity of NIs. Few studies have examined the effect of limonene, MBOA and MHPA on ammonia oxidizer isolates, focusing solely on *Nitrosomonas europaea* (Ward et al., 1997; Gopalakrishnan et al., 2007; Otaka et al., 2023) and thus, an evaluation of these BNIs on ammonia oxidizer isolates is required to further understand BNIs specificity and mode of action. In soil, as expected, different ammonia oxidizer groups had different responses to the NI treatments in the AS and CS. Yet, differences in *amoA* gene and transcripts numbers, as well as the presence or absence of community composition shifts, were not always consistent within a group of ammonia oxidizers, or across soils. For example, although the significant inhibitory effect of DMPP on AOB was evident in both *amoA* gene and transcript copy numbers (Fig. S2 Fig. 2), no such effect was observed on AOA communities. Similarly, and as previously shown (Bachtsevani et al., 2021; Papadopoulou et al., 2024), the relative abundance of members of the terrestrial AOB genus *Nitrosospira* were significantly reduced by the SNI DMPP in the AS (Fig. 3A). In contrast, AOA *amoA* transcript abundance remained significantly higher in DMPP microcosms compared to the BNI or control microcosms from both soils (Fig 2A), but this was not mirrored in the AOA *amoA* gene abundance or community composition, which remained relatively stable (Fig S2, S3, S4). This mild reduction in AOA *amoA* transcript abundances is likely a response to the concomitant rise in AOB *amoA* transcript levels and transcript-to-gene ratios (Fig. 2). A similar dynamic, likely driven by niche differentiation or competition between ammonia oxidizer populations, has also been observed in other studies (Hink et al., 2018; Papadopoulou et al., 2024). Notably, the effect of BNIs on ammonia oxidizer communities, except for limonene, was overall low, regardless of whether *amoA* gene abundance, transcript abundance, or community composition was used to assess the effect. These results are contrary to previous studies, which have reported a higher sensitivity of AOA over AOB to BNIs (Kaur-Bhambra et al., 2022).

Interestingly, although NO_X_^-^ accumulation in the AS was reduced by MBOA, AOB *amoA* gene and transcript abundances remained high and increased towards the end of the incubation period, which could be a response mechanism of AOB to overcome inhibition. A similar compensatory mechanism has been observed in pure culture studies through *amoA* upregulation under ammonia starvation or during low energy availability (Bollmann et al., 2005; Nakagawa and Stahl, 2013). This response mechanism serves as a strategy to ensure fast recovery once ammonia becomes available again or, in this case, once inhibition is relieved. Consequently, an increase in *amoA* transcript abundance does not necessarily indicate a lack of effect from the BNI but might be a response to energy limitation. In summation, the response of ammonia-oxidizing communities to BNIs is shaped by microbial physiology and soil-specific community composition, highlighting the importance of a deeper understanding of the mode of action of specific BNIs on key soil ammonia oxidizers, as well as a cautious interpretation of gene and transcript abundance data.

### 4.3. Response of non-target soil microorganisms and fungi to NI application

BNIs are often carbon-rich compounds, some with allelopathic functions (Bachheti et al., 2020), and it is possible that their application in soil will introduce changes in the abundance and composition of the native microbial community. Many of these natural compounds are also involved in plant signaling and defense (Subbarao et al., 2006; Nardi et al., 2020), making it essential to understand BNIs effects on non-nitrifying community members. In fact, previous studies have observed that SNI application also affects microbial community members other than ammonia oxidizers (Bachtsevani et al., 2021; Papadopoulou et al., 2022; Ramotowski and Shi, 2022). We therefore hypothesized that both microbial and fungal communities will be affected by the application of limonene, MHPA, and MBOA. Accordingly, BNI application stimulated a greater number of non-target bacterial and fungal taxa than it suppressed in both soils (Fig. 3). Despite showing a low NI efficacy and close to no effect on ammonia oxidizers (Fig. 1, 2, S2-S4), MHPA significantly increased the relative abundance of several bacterial genera (Fig. 3). Microbial degradation of MHPA by soil bacteria has been previously reported (Hirakawa et al., 2012), suggesting that the low efficacy observed here may be due to MHPA utilization as a carbon source by heterotrophic soil microorganisms. Importantly, MHPA application led to an increased relative abundance of *Fusarium* in the AS, a genus that includes some plant pathogens (Fig. 3B). The release from bacterial and fungal competitors along with changes in nutrient availability — such as shifts in nitrogen form or the release of BNI degradation products — could create favorable conditions for fast-growing fungi to colonize the soil matrix. Despite being the least effective BNI, this result highlights the importance of identifying the overall effects of NI application in soil beyond nitrification, as NI-induced changes could result in indirect and undesired effects for crop health.

Limonene strongly suppressed NOx-formation, stimulated NH₄⁺ immobilization (Fig. 1), and had a strong impact on ammonia oxidizer communities (Fig. 2, S2-S4), all of which are likely an indirect effect driven by its impact on the non-nitrifying microbial communities. The marked increase in the relative abundance of hydrocarbon degraders, with *Rhodococcus*-related ASVs comprising 50% and 30% of the bacterial community in the AS and CS at the end of the incubation period, respectively (Fig. 3) is likely due to the utilization of limonene as a carbon and/or energy source (Mirata et al., 2009; Marmulla and Harder, 2014). Furthermore, limonene also induced a pronounced increase in the relative abundance of the genus *Fusicolla*, suggesting this taxon likely contributed to the high NH₄⁺ immobilization observed under limonene treatment in both soils. DMPP demonstrated high efficacy in reducing NOx⁻ accumulation and showed no off-target effects on microbial and minimal off-target effects on fungal communities in both soils. However, DMPP stimulated NH₄⁺ immobilization in the CS, and other off -target effects have been reported before (Maienza et al., 2014; Bachtsevani et al., 2021; Liu et al., 2024). Notably, MBOA also had minimal effects on the microbial and fungal communities in both soils, as it only increased the relative abundance of a methylotroph that, similar to what was observed with the other BNIs, likely uses it as a carbon source (Macey et al., 2018). This, combined with its relatively high efficacy in inhibiting NOx⁻ formation, and NH_4_^+^ retention, makes MBOA a promising BNI candidate.

### 4.4. Persistence of recoverable NI fraction in soil

NI persistence and stability in soil are key factors when it comes to their application in agricultural soils. Previous studies have shown that the fate of NIs in soils is determined by soil physicochemical properties, NI type, application rate, as well as the soil-specific microbial community (Barth et al., 2008; McGeough et al., 2016; Guardia et al., 2018; Ma et al., 2021). Accordingly, differences in BNI persistence across the AS and CS were observed, along with an overall lower persistence of BNIs compared to the SNI DMPP (Fig. 4). Higher organic matter as well as clay content are usually associated with higher soil NI sorption, which in turn affects NI bioavailability and efficacy (Subbarao et al., 2006; Guardia et al., 2018; Nardi et al., 2020). Compared to the CS, the AS not only has significantly higher pH, but also a higher cation exchange capacity, CaCO_3_ and total C content (Rojas-Pinzon et al., 2024), which all contribute to a higher NI sorption capacity. Yet, the lowest NI recoveries were observed in the CS (Table S11). Although higher sorption was expected in the AS, the lower NIs recovery in the CS is likely due to the low application rates of some BNIs (between 2-3 times less limonene and MBOA were applied in the CS), which may have also resulted in the low BNI efficacy observed in this soil. These concentrations were previously estimated to cause ∼80% inhibition (Rojas-Pinzon et al., 2024), which highlights both the effect of the experimental setup (e.g., slurry vs. microcosm incubations) as well as of the application rate in determining NI efficacy in soil.

As in other studies, where the dissipation time of DMPP has been estimated to be between 5-28 days (Doran et al., 2018), DMPP was the most persistent NI tested here, with >40% recovery after 15 and 21 days in the AS and the CS, respectively. Among the NIs tested, limonene showed the lowest persistence in both soils (Fig. 4). The limonene-induced blooms of hydrocarbon degraders likely explain both its low persistence as well as the substantial drop in inorganic N (Figure 1C) observed after its application. Stimulation of NH4⁺ immobilization after limonene application has been previously observed and attributed to the carbon input that limonene provides to the soil microbial community (Bremner and McCarty, 1988). MHPA showed both low efficacy (Fig. 1B) and low persistence (Fig. 4). Phenylpropanoids such as MHPA are known to act as substrates for the soil microbial community, but also as toxins, and thus mediate plant defense mechanisms (Li et al., 2018; Zwetsloot et al., 2020). Since MHPA stimulated more bacterial and fungal genera than it suppressed (Fig. 3), it likely served as a carbon source to diverse community members.

Interestingly, MBOA showed a relatively high recovery rate and longer persistence time (36.3% recovery after 15 days of application in the AS), but it loses its efficacy as NI after 8 days. Microbial degradation thus likely led to a reduction of MBOA concentration below an inhibitory threshold. A previous study also observed microbial MBOA degradation, as MBOA concentrations declined within 4–5 days in non-sterilized soil but remained stable in sterilized controls (Otaka et al., 2023). Thus, despite showing only limited off-target effects (Fig. 3), microbial degradation plays a key role for MBOA efficacy in soil and needs to be further examined to determine suitable application rates. Overall, a better understanding of the fate of BNIs in soil is essential for designing effective application strategies that ensure sufficient persistence to inhibit nitrification, while avoiding prolonged activity that could result in unintended effects on non-target members of the microbial community.

## 5. Conclusion

Our study provides additional understanding on the effect of three BNI compounds (limonene, MHPA, and MBOA) on target and non-target microbial communities, as well as on their fate in two contrasting agricultural soils. We showed that MBOA may be a suitable biological nitrification inhibitor, as it displayed comparatively high efficacy combined with low off-targeted effects. However, a better understanding of MBOA degradation across soils is needed to design adequate application strategies. We also showed that although limonene effectively inhibits NOx⁻ accumulation and could therefore be considered a BNI, off-target shifts in bacterial and fungal community composition likely stimulated NH4⁺ immobilization and reduced NOx⁻ accumulation. In summary, our results highlight that evaluating BNI efficacy in soil requires a more complex experimental framework than is commonly used in many studies proposing novel inhibitory compounds. Overall, the identification of the specific mode of action of BNIs on nitrifiers, the off-target microbial taxa affected that could also potentially influence BNI stability in soil, as well as the factors determining BNI availability and persistence should all be considered when it comes to finding natural and sustainable alternatives to reduce the current environmental impact of fertilization.

## Acknowledgements

We thank the Austrian Agency for Health and Food Safety (AGES) for facilitating the sampling sites. We also thank the Institute of Bioanalytics and Agro-Metabolomics of the University of Natural Resources and Life Sciences for facilitating the quantification of the nitrification inhibitor extracts. We are grateful to Julia Ramesmayer and Jasmin Schwarz for their technical support with the sequencing of the samples, as well as Joana Seneca for the pre-processing of the sequencing datasets.

## CRediT authorship contribution statement

**Paula A. Rojas-Pinzon:** Conceptualization, Data curation, Formal analysis, Investigation, Methodology, Visualization, Writing – original draft, Writing – review & editing. **Bernhard Seidl:** Methodology, Data curation. **Stella Kejik:** Methodology. **Christopher J. Sedlacek:** Conceptualization, Funding acquisition, Project administration, Supervision, Writing –original draft, Writing –review & editing. **Judith Prommer:** Methodology. **Christoph Bueschl:** Funding acquisition, Writing – review & editing. **Taru Sanden:** Resources. **Heide Spiegel:** Resources. **Andrew T. Giguere:** Conceptualization, Funding acquisition, Project administration, Supervision, Writing – review & editing. **Lucia Fuchslueger:** Conceptualization, Funding acquisition, Project administration, Supervision, Writing –review & editing. **Petra Pjevac:** Conceptualization, Funding acquisition, Project administration, Supervision, Writing – original draft, Writing – review & editing.

## Declaration of Competing Interest

The authors declare no conflict of interest or competing interests.

## Data availability

Amplicon sequence data was deposited at the NCBI Sequence Read Archive (SRA) under accession number PRJNA1312475

## Funding

This research was funded by the Austrian Science Fund (FWF) Young Investigators Research Grant program (grant number ZK-74B, DOI: 10.55776/ZK74) granted to (P.P., C.J.S, C.B., A.T.G., and L.F.).

## Notes

### Competing Interest Statement

The authors have declared no competing interest.

