## Supplementary figures and tables for "Beyond efficacy: Persistence and off-target effects of three biological nitrification inhibitors in two contrasting agricultural soils"

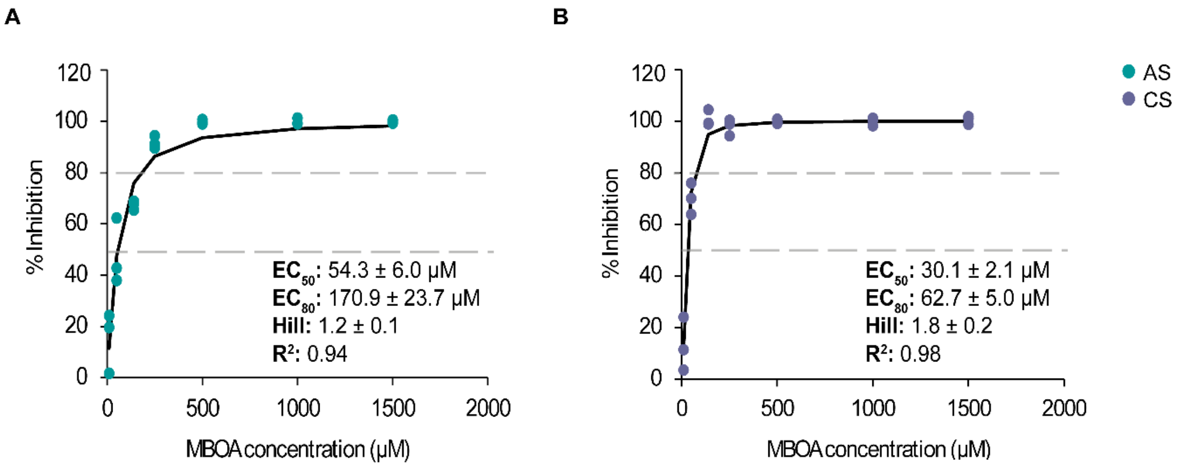


**Fig. S1.** Log-logistic model fitting to estimate the $EC_{50}$ and $EC_{80}$ values of MBOA in the (A) AS and (B) CS. The $EC_{50}$ , $EC_{80}$ values (dashed lines), hill coefficient, as well as the $R^2$ with their respective standard errors are displayed. For further information about the methods and calculations, refer to Rojas et al. (2024).

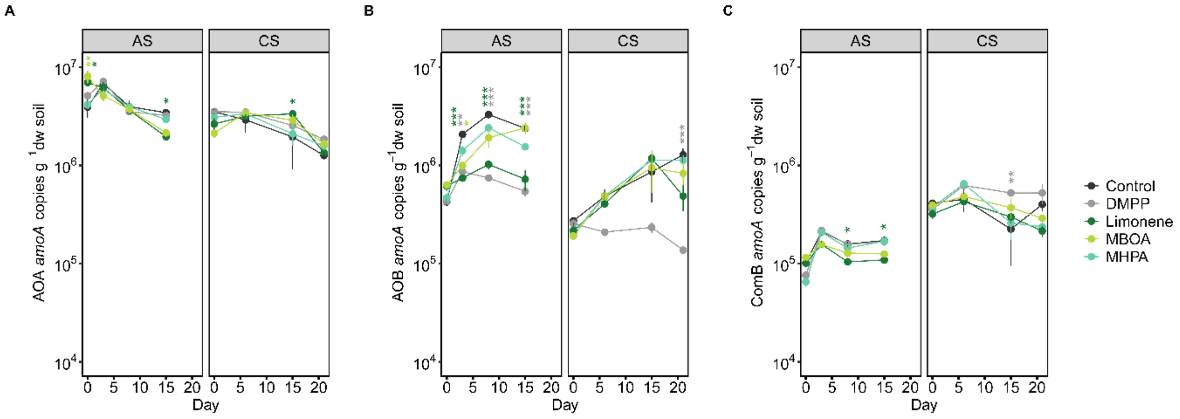


**Fig. S2.** Change in *amoA* gene abundance of the (A) AOA, (B) AOB, and (C) ComB communities in the AS and CS with and without addition of NIs. Data are normalized per gram dry weight (dw) soil and presented as means ± SE (n = 4). Asterisks and the color denote significant differences of a specific treatment relative to the control on each day (* < 0.05, ** < 0.01, *** < 0.001). The effects of the NI treatment and incubation time were tested using a two-way ANOVA followed by a Tukey HSD for data with normal distribution and homogeneous variances (AOA *amoA* abundances at the AS). When these assumptions were not met, a generalized lineal model with pairwise comparisons of estimated marginal means (EMMs) (AOA and ComB *amoA* abundances at the CS, and AOB *amoA* abundances of both sites) or a generalized least squares model with EMMs pairwise comparisons (ComB *amoA* abundances at the AS) were used instead.

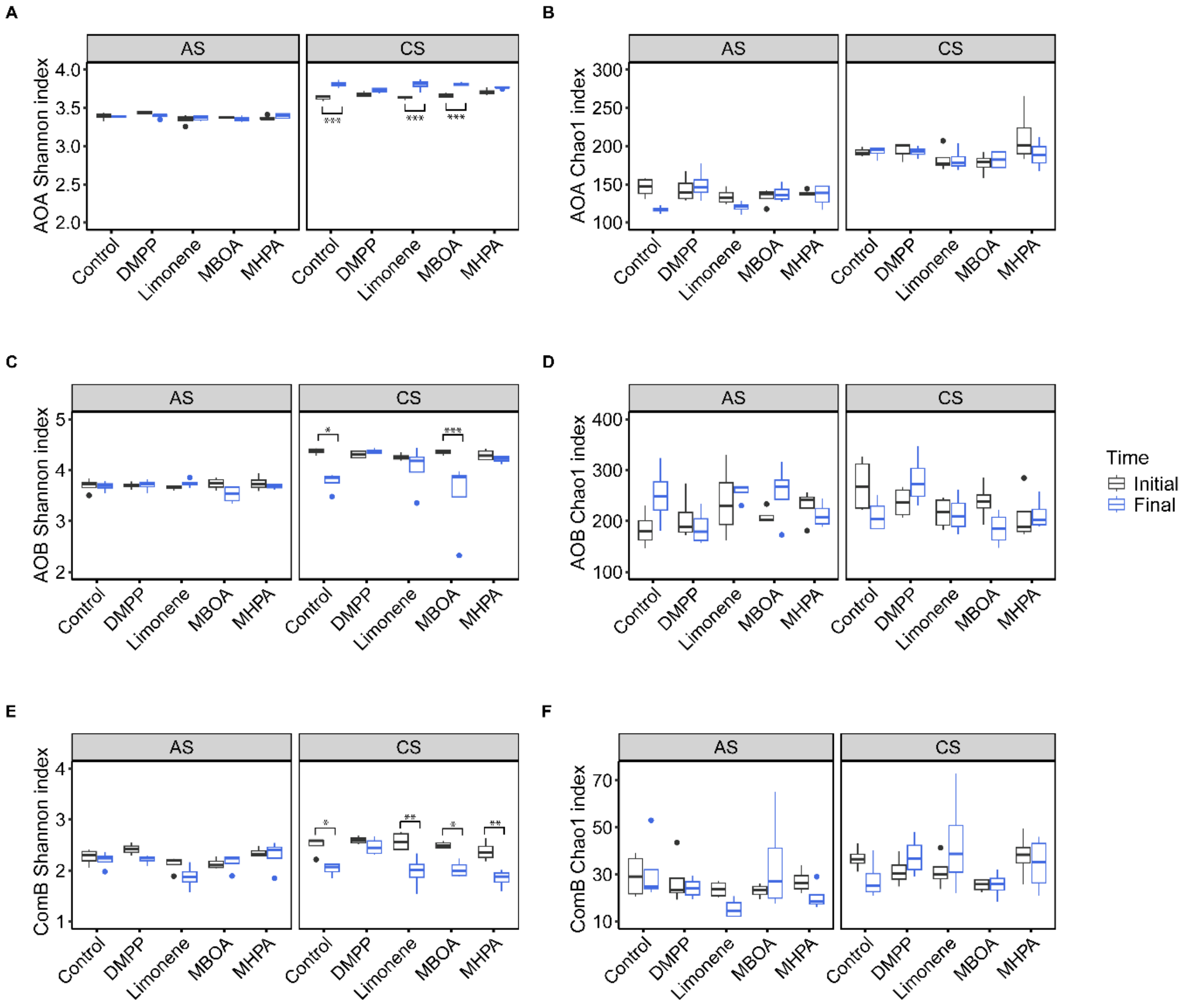


**Fig. S3.** α-diversity indexes of the *amoA* gene-based community composition of the AS and CS from the initial and final timepoint (15 days for the AS and 21 days for the CS) of the incubation. The Shannon diversity index and Chao1 index for the AOA *amoA* communities (A and B), AOB *amoA* communities (C and D), and ComB *amoA* communities (E and F) are shown. The median, first and third quartile of each diversity index are depicted as the middle, upper, and lower hinges, respectively in the boxplots. The length of the whiskers is determined by the largest and the smallest value in the dataset that are within 1.5 times the interquartile range and outliers are shown as points. Asterisks denote significant differences between time points within each treatment (* < 0.05, ** < 0.01, *** < 0.001).

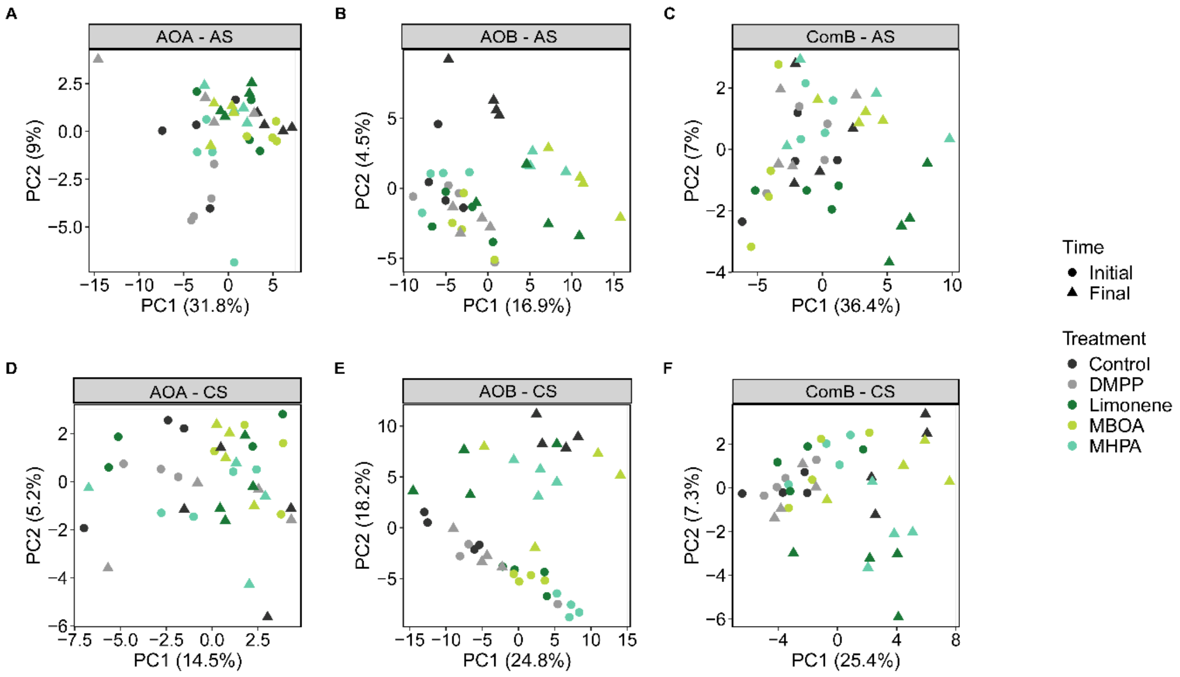


**Fig. S4.** Effect of NIs on the ammonia oxidizer community β-diversity in the AS and the CS at the initial and final timepoint (15 days for the AS and 21 days for the CS) of the incubation. Principal component analysis at the ASV-level on an Aitchison distance matrix for the (A-C) AOA, AOB and ComB *amoA* in the AS, and for the (D-F) AOA, AOB and ComB *amoA* in the CS.

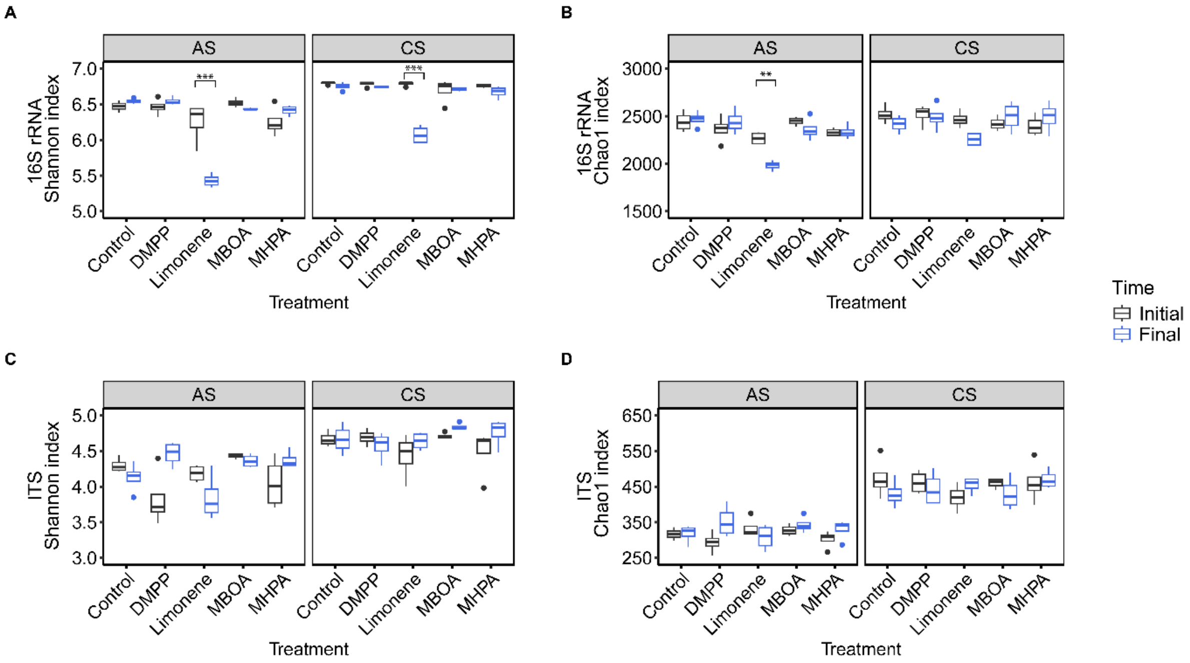


**Fig. S5.** α-diversity indexes of the soil bacterial and fungal community composition of the AS and CS at the initial and final timepoint (15 days for the AS and 21 days for the CS) of the incubation. The Shannon diversity index and Chao1 index for the 16S rRNA gene-based communities (A and B), and for the ITS2 gene-based communities (C and D) are shown. The median, first and third quartile of each diversity index are depicted as the middle, upper, and lower hinges, respectively in the boxplots. The length of the whiskers is determined by the largest and the smallest value in the dataset that are within 1.5 times the interquartile range and outliers are depicted by points. Asterisks denote significant differences between time points within each treatment (* < 0.05, ** < 0.01, *** < 0.001).

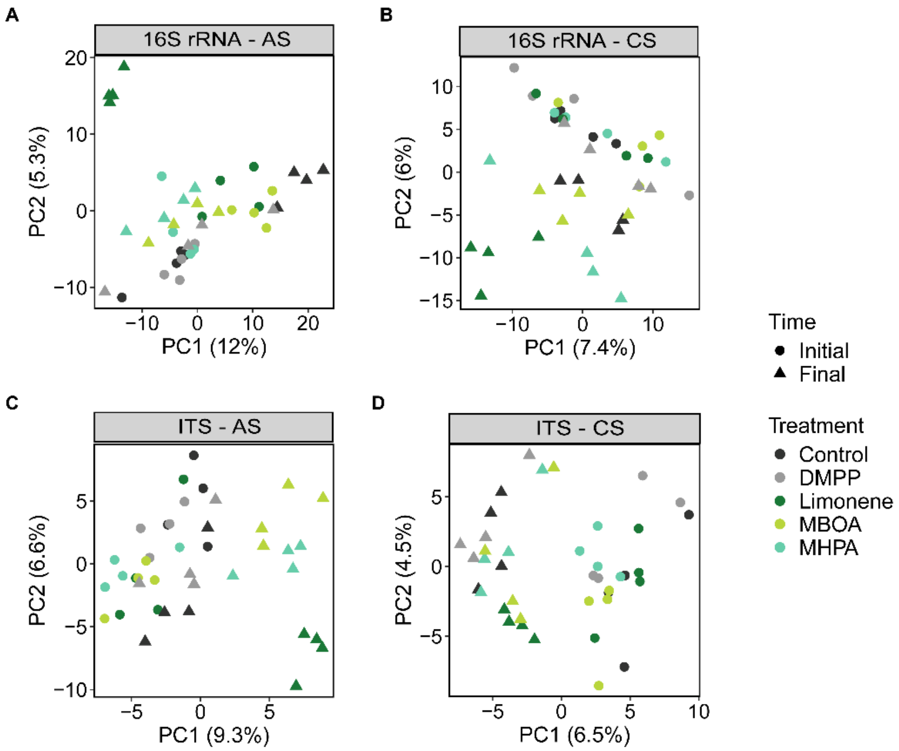


**Fig. S6.** Effect of NI on the β-diversity of the bacterial and fungal communities of the AS and the CS at the initial and final timepoint (15 days for the AS and 21 days for the CS) of the incubation. Principal component analysis at the ASV-level on an Aitchison distance matrix for the 16S rRNA gene-based communities in the (A) AS and (B) in the CS; and for the ITS2 gene-based communities in the (C) AS and (D) in the CS.

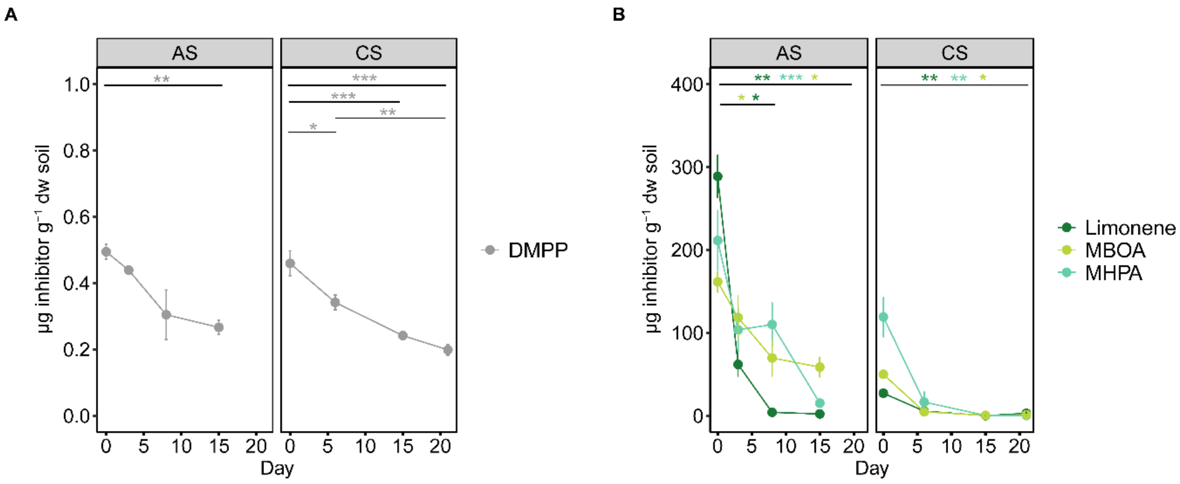


**Fig. S7.** Change in concentration throughout time for (A) the SNI and (B) the BNIs. Data are normalized per gram dry weight (dw) soil and presented as means ± SE (n = 4). Asterisks and the color denote significant differences in the concentration of each NI (*< 0.05, **< 0.01, *** < 0.001).

**Table S1**: The primers and conditions used for qPCR, RT-qPCR, and amplicon sequencing in this study.

| Gene | Primer name | qPCR/PCR conditions | Primer final concentration | Reference |
|---|---|---|---|---|
| AOA *amoA* | 104F<br>616R | 95°C for 15 min<br>40 cycles of:<br>95°C for 15 s<br>60°C for 45 s<br>78°C for 10 s<br>Final extension:<br>60°C for 10 min<br>Melting curve starting at 55°C | 1 µM | Alves et al. 2013 |
| AOB *amoA* | 1F<br>2R | 94°C for 5 min<br>40 cycles of:<br>94°C for 1 min<br>60°C for 1 min 30 s<br>72°C for 1 min 30 s<br>Melting curve starting at 45°C | 0.3 µM | Rotthauwe et al., 1997 |
| ComB *amoA* | comaB_244F<br>comaB_659R | 95°C for 3 min<br>40 cycles of:<br>95°C for 30 s<br>52°C for 45 s<br>72°C for 1 min<br>Melting curve starting at 38°C | 0.5 µM | Pjevac et al., 2017 |
| ComA *amoA* | comaA_244F<br>comaA_659R | 95°C for 3 min<br>40 cycles of:<br>95°C for 30 s<br>52°C for 45 s<br>72°C for 1 min<br>Melting curve starting at 38°C | 0.5 µM | Pjevac et al., 2017 |
| 16S rRNA | 515F<br>806R | 95°C for 2 min<br>30 cycles of:<br>95°C for 20 s<br>55°C for 15 s<br>72°C for 5 min<br>72°C for 10 min | 0.2 µM | Apprill et al., 2015; Parada et al., 2016 |
| ITS2 | gITS7<br>ITS4 | 94°C for 5 min<br>30 cycles of:<br>94°C for 30 s<br>56°C for 30 s<br>72°C for 30s<br>72 for 7 min | 0.2 µM | Ihrmark et al., 2012 |

**Table S2**: Remaining $NH_4^+$ on the final day of the incubation with and without NIs for the AS (day 15) and the CS (day 21). Data are presented as means ± SE (n = 4). For each treatment, the initial $NH_4^+$ concentrations at day 0 were taken as 100%.

| | AS | | | CS | | |
|---|---|---|---|---|---|---|
| Treatment | Mean (µg $NH_4^+$ $g^{-1}$ soil) | SE | Fraction of remaining $NH_4^+$ | Mean (µg $NH_4^+$ $g^{-1}$ soil) | SE | Fraction of remaining $NH_4^+$ |
| Control | 0.6 | ± 0.2 | 0.4% | 27.1 | ± 2.6 | 21.4% |
| DMPP | 58.3 | ± 2.8 | 46.0% | 73.4 | ± 0.6 | 58.1% |
| Limonene | 26.2 | ± 4.3 | 20.7% | 32.6 | ± 11.5 | 26.0% |
| MBOA | 26.4 | ± 5.2 | 21.0% | 54.1 | ± 7.1 | 43.0% |
| MHPA | 7.1 | ± 5.5 | 5.6% | 39.0 | ± 2.3 | 30.7% |

**Table S3**: Initial and final qPCR absolute values and fold changes for the AOA, AOB and ComB *amoA* genes in the AS and the CS (A). T-test or the non-parametric Mann-Whitney test were done to test for significant differences between the initial and final *amoA* gene abundances within each treatment. (B) The t-value (for t-test), U-statistic (for the Mann-Whitney test) as well as the p-value are depicted. Data are presented as means ± SE (n = 4) and significant differences are shown in bold.

A

| | | AS | | | CS | | |
|---|---|---|---|---|---|---|---|
| Gene | Treatment | Initial *amoA* copies ± SE | Final *amoA* copies ± SE | Fold change | Initial *amoA* copies ± SE | Final *amoA* copies ± SE | Fold change |
| AOA *amoA* | Control | $3.9 \times 10^6 \pm 8.6 \times 10^5$ | $3.4 \times 10^6 \pm 1.2 \times 10^5$ | 0.9 | $3.5 \times 10^6 \pm 2.4 \times 10^5$ | $1.3 \times 10^6 \pm 1.2 \times 10^5$ | 0.4 |
| | DMPP | $5.1 \times 10^6 \pm 4.6 \times 10^5$ | $3.2 \times 10^6 \pm 2.3 \times 10^6$ | 0.6 | $3.5 \times 10^6 \pm 1.1 \times 10^5$ | $1.8 \times 10^6 \pm 1.9 \times 10^5$ | 0.5 |
| | Limonene | $7.0 \times 10^6 \pm 5.7 \times 10^5$ | $2.0 \times 10^6 \pm 1.4 \times 10^5$ | 0.3 | $2.6 \times 10^6 \pm 3.7 \times 10^5$ | $1.35 \times 10^6 \pm 4.1 \times 10^4$ | 0.5 |
| | MBOA | $8.0 \times 10^6 \pm 1.2 \times 10^6$ | $2.1 \times 10^6 \pm 2.1 \times 10^5$ | 0.3 | $2.1 \times 10^6 \pm 2.1 \times 10^5$ | $1.7 \times 10^6 \pm 2.0 \times 10^5$ | 0.8 |
| | MHPA | $4.1 \times 10^6 \pm 3.8 \times 10^5$ | $2.9 \times 10^6 \pm 2.2 \times 10^5$ | 0.7 | $3.1 \times 10^6 \pm 4.7 \times 10^5$ | $1.6 \times 10^6 \pm 1.7 \times 10^5$ | 0.5 |
| AOB *amoA* | Control | $4.2 \times 10^5 \pm 3.9 \times 10^4$ | $2.4 \times 10^6 \pm 1.4 \times 10^5$ | 5.6 | $2.7 \times 10^5 \pm 8.7 \times 10^3$ | $1.3 \times 10^6 \pm 1.9 \times 10^5$ | 4.7 |
| | DMPP | $4.3 \times 10^5 \pm 2.2 \times 10^4$ | $5.4 \times 10^5 \pm 5.9 \times 10^4$ | 1.3 | $2.6 \times 10^5 \pm 2.9 \times 10^4$ | $1.4 \times 10^5 \pm 9.9 \times 10^3$ | 0.5 |
| | Limonene | $6.2 \times 10^5 \pm 6.8 \times 10^4$ | $7.2 \times 10^5 \pm 1.7 \times 10^5$ | 1.2 | $2.2 \times 10^5 \pm 4.4 \times 10^4$ | $4.9 \times 10^5 \pm 1.4 \times 10^5$ | 2.2 |
| | MBOA | $6.3 \times 10^5 \pm 5.7 \times 10^4$ | $2.4 \times 10^6 \pm 3.5 \times 10^5$ | 3.8 | $1.9 \times 10^5 \pm 1.7 \times 10^4$ | $8.3 \times 10^5 \pm 2.0 \times 10^5$ | 4.3 |
| | MHPA | $4.7 \times 10^5 \pm 4.5 \times 10^4$ | $1.5 \times 10^6 \pm 1.3 \times 10^5$ | 3.3 | $2.0 \times 10^5 \pm 1.7 \times 10^4$ | $1.1 \times 10^6 \pm 1.4 \times 10^5$ | 5.5 |
| ComB *amoA* | Control | $1.0 \times 10^5 \pm 6.6 \times 10^3$ | $1.7 \times 10^5 \pm 1.7 \times 10^4$ | 1.7 | $4.1 \times 10^5 \pm 2.9 \times 10^4$ | $4.0 \times 10^5 \pm 6.2 \times 10^4$ | 0.9 |
| | DMPP | $7.6 \times 10^4 \pm 8.1 \times 10^3$ | $1.7 \times 10^5 \pm 1.8 \times 10^4$ | 2.2 | $3.5 \times 10^5 \pm 2.3 \times 10^4$ | $5.2 \times 10^5 \pm 1.2 \times 10^5$ | 1.5 |
| | Limonene | $1.0 \times 10^5 \pm 7.8 \times 10^3$ | $1.1 \times 10^5 \pm 5.1 \times 10^3$ | 1.1 | $3.2 \times 10^5 \pm 3.4 \times 10^4$ | $2.1 \times 10^5 \pm 2.8 \times 10^4$ | 0.7 |
| | MBOA | $1.2 \times 10^5 \pm 7.5 \times 10^3$ | $1.3 \times 10^5 \pm 7.9 \times 10^3$ | 1.1 | $3.9 \times 10^5 \pm 3.8 \times 10^4$ | $2.9 \times 10^5 \pm 1.8 \times 10^4$ | 0.7 |
| | MHPA | $6.5 \times 10^4 \pm 7.4 \times 10^3$ | $1.7 \times 10^5 \pm 1.4 \times 10^4$ | 2.6 | $3.7 \times 10^5 \pm 2.7 \times 10^4$ | $2.4 \times 10^5 \pm 4.5 \times 10^4$ | 0.6 |

B

| Gene | Treatment | AS | | CS | |
|---|---|---|---|---|---|
| | | t-value / U-statistic† | p-value | t-value / U-statistic† | p-value |
| AOA *amoA* | Control | 0.55 | 0.621 | **8.40** | **< 0.001** |
| | DMPP | **3.67** | **0.010** | **7.63** | **< 0.001** |
| | Limonene | **8.57** | **0.002** | **3.44** | **0.014** |
| | MBOA | **6.39** | **< 0.001** | 1.59 | 0.162 |
| | MHPA | **16†** | **0.029** | **3.02** | **0.023** |
| AOB *amoA* | Control | **0†** | **0.029** | **0†** | **0.028** |
| | DMPP | -1.86 | 0.111 | **3.87** | **0.008** |
| | Limonene | -0.57 | 0.585 | -1.77 | 0.126 |
| | MBOA | **-5.01** | **0.002** | **-4.49** | **0.004** |
| | MHPA | **-7.99** | **< 0.001** | **-6.43** | **< 0.001** |
| ComB *amoA* | Control | **-3.87** | **0.008** | **-3.86** | **0.008** |
| | DMPP | **-4.69** | **0.003** | **-4.68** | **0.003** |
| | Limonene | -0.85 | 0.427 | 2.35 | 0.056 |
| | MBOA | -0.91 | 0.397 | **2.50** | **0.046** |
| | MHPA | **-6.60** | **< 0.001** | **2.59** | **0.040** |

†Cases where the non-parametric Mann-Whitney test was used

**Table S4**: Post-hoc test results for $NH_4^+$ consumption differences between treatments on each day for (A) the AS and (B) the CS. For the AS, Tukey HSD comparisons were done after a two-way ANOVA, while pairwise comparisons of estimated marginal means (EMMs) from a gamma-log generalized linear model (glm) were used for the CS. Significant differences relative to the control and between treatments are shown in bold.

A

| **Comparison** | **Day 0** | | **Day 2** | | **Day 3** | | **Day 8** | | **Day 15** | |
|---|---|---|---|---|---|---|---|---|---|---|
| | **diff** | **p adj** | **diff** | **p adj** | **diff** | **p adj** | **diff** | **p adj** | **diff** | **p adj** |
| DMPP-Control | -20.5 | 0.102 | 10.0 | 0.989 | -3.0 | 1 | **32.5** | **< 0.001** | **57.8** | **0** |
| Limonene-Control | -9.2 | 0.996 | **25.5** | **0.007** | 13.8 | 0.766 | 11.3 | 0.953 | **25.7** | **0.007** |
| MBOA-Control | -15.2 | 0.596 | 20.4 | 0.105 | 8.8 | 0.997 | **41.4** | **< 0.001** | **25.8** | **< 0.001** |
| MHPA-Control | -20.1 | 0.121 | 2.2 | 1 | -3.4 | 1 | 11.0 | 0.965 | 6.5 | 0.999 |
| Limonene-DMPP | 11.3 | 0.954 | 15.5 | 0.559 | 16.8 | 0.398 | -21.1 | 0.074 | **-32.1** | **< 0.001** |
| MBOA-DMPP | 5.2 | 0.999 | 10.4 | 0.981 | 11.8 | 0.929 | 8.8 | 0.997 | **-31.9** | **< 0.001** |
| MHPA-DMPP | 0.4 | 1 | -7.7 | 0.999 | -0.4 | 1 | -21.5 | 0.064 | **-51.2** | **0** |
| MBOA-Limonene | -6.1 | 0.999 | -5.1 | 0.999 | -4.9 | 0.999 | **30.0** | **< 0.001** | 0.1 | 1 |
| MHPA-Limonene | -11.0 | 0.967 | **-23.2** | **0.026** | -17.2 | 0.350 | -0.3 | 1 | -19.1 | 0.180 |
| MHPA-MBOA | -4.9 | 0.999 | -18.2 | 0.259 | -12.3 | 0.903 | **-30.4** | **< 0.001** | -19.2 | 0.170 |

B

| **Comparison** | **Day 0** | | **Day 3** | | **Day 6** | | **Day 15** | | **Day 21** | |
|---|---|---|---|---|---|---|---|---|---|---|
| | **Est.** | **p adj** | **Est.** | **p adj** | **Est.** | **p adj** | **Est.** | **p adj** | **Est.** | **p adj** |
| DMPP-Control | 0.039 | 1 | 0.012 | 1 | -0.112 | 1 | -0.247 | 1 | **-0.997** | **< 0.001** |
| Limonene-Control | 0.057 | 1 | 0.081 | 1 | 0.187 | 1 | 0.207 | 1 | -0.186 | 1 |
| MBOA-Control | -0.027 | 1 | 0.010 | 1 | -0.134 | 1 | -0.215 | 1 | **-0.691** | **< 0.001** |
| MHPA-Control | 0.039 | 1 | 0.048 | 1 | -0.072 | 1 | -0.065 | 1 | -0.365 | 1 |
| Limonene-DMPP | 0.017 | 1 | 0.068 | 1 | 0.299 | 1 | 0.454 | 0.106 | **0.810** | **< 0.001** |
| MBOA-DMPP | -0.067 | 1 | -0.002 | 1 | -0.021 | 1 | 0.031 | 1 | 0.305 | 1 |
| MHPA-DMPP | 0.000 | 1 | 0.035 | 1 | 0.040 | 1 | 0.181 | 1 | **0.631** | **< 0.001** |
| MBOA-Limonene | -0.084 | 1 | -0.071 | 1 | -0.321 | 1 | -0.422 | 0.250 | **-0.505** | **0.025** |
| MHPA-Limonene | -0.017 | 1 | -0.033 | 1 | -0.259 | 1 | -0.272 | 1 | -0.178 | 1 |
| MHPA-MBOA | 0.067 | 1 | 0.037 | 1 | 0.061 | 1 | 0.149 | 1 | 0.326 | 1 |

**Table S5**: Post-hoc test results for $NO_x^-$ ($NO_3^-$ and $NO_2^-$) accumulation differences between treatments on each day for (A) the AS and (B) the CS. Pairwise comparisons of estimated marginal means (EMMs) from a gamma-log generalized linear model (glm) and an inverse gaussian glm were used for the AS and CS, respectively. Significant differences relative to the control as well as between treatments are depicted in bold.

A

| Comparison | Day 0 | | Day 2 | | Day 3 | | Day 8 | | Day 15 | |
|---|---|---|---|---|---|---|---|---|---|---|
| | Est. | p adj | Est. | p adj | Est. | p adj | Est. | p adj | Est. | p adj |
| DMPP-Control | -0.299 | 1 | 0.218 | 1 | 0.525 | 0.112 | **0.714** | **< 0.001** | **0.758** | **< 0.001** |
| Limonene-Control | -0.323 | 1 | **0.952** | **< 0.001** | **1.333** | **< 0.001** | **1.455** | **< 0.001** | **1.271** | **< 0.001** |
| MBOA-Control | -0.453 | 0.571 | **0.597** | **0.019** | **0.689** | **0.002** | **0.844** | **< 0.001** | 0.148 | 1 |
| MHPA-Control | -0.293 | 1 | 0.144 | 1 | 0.375 | 1 | 0.315 | 1 | 0.128 | 1 |
| Limonene-DMPP | -0.024 | 1 | **0.743** | **< 0.001** | **0.808** | **< 0.001** | **0.740** | **< 0.001** | 0.512 | 0.150 |
| MBOA-DMPP | -0.154 | 1 | 0.378 | 1 | 0.164 | 1 | 0.129 | 1 | **-0.609** | **0.013** |
| MHPA-DMPP | 0.006 | 1 | -0.073 | 1 | -0.149 | 1 | -0.398 | 1 | **-0.629** | **0.008** |
| MBOA-Limonene | -0.130 | 1 | -0.355 | 1 | **-0.644** | **0.005** | **-0.610** | **0.013** | **-1.122** | **< 0.001** |
| MHPA-Limonene | 0.030 | 1 | **-0.807** | **< 0.001** | **-0.958** | **< 0.001** | **-1.139** | **< 0.001** | **-1.142** | **< 0.001** |
| MHPA-MBOA | 0.160 | 1 | -0.452 | 0.590 | -0.313 | 1 | -0.528 | 0.104 | -0.019 | 1 |

B

| Comparison | Day 0 | | Day 3 | | Day 6 | | Day 15 | | Day 21 | |
|---|---|---|---|---|---|---|---|---|---|---|
| | Est. | p adj | Est. | p adj | Est. | p adj | Est. | p adj | Est. | p adj |
| DMPP-Control | 0.079 | 1 | 0.100 | 1 | 0.382 | 0.850 | **0.871** | **< 0.001** | **1.160** | **< 0.001** |
| Limonene-Control | 0.329 | 0.094 | 0.320 | 0.856 | **0.624** | **< 0.001** | 0.370 | 1 | 0.507 | 1 |
| MBOA-Control | 0.125 | 1 | 0.082 | 1 | 0.386 | 0.747 | 0.558 | 0.675 | 0.303 | 1 |
| MHPA-Control | 0.145 | 1 | 0.162 | 1 | 0.296 | 1 | 0.300 | 1 | 0.165 | 1 |
| Limonene-DMPP | 0.249 | 1 | 0.219 | 1 | 0.242 | 1 | -0.500 | 0.336 | **-0.652** | **0.019** |
| MBOA-DMPP | 0.046 | 1 | -0.017 | 1 | 0.005 | 1 | -0.312 | 1 | **-0.857** | **< 0.001** |
| MHPA-DMPP | 0.066 | 1 | 0.062 | 1 | -0.086 | 1 | -0.570 | 0.082 | **-0.994** | **< 0.001** |
| MBOA-Limonene | -0.204 | 1 | -0.237 | 1 | -0.237 | 1 | 0.187 | 1 | -0.204 | 1 |
| MHPA-Limonene | -0.183 | 1 | -0.157 | 1 | -0.327 | 0.999 | -0.070 | 1 | -0.342 | 1 |
| MHPA-MBOA | 0.020 | 1 | 0.080 | 1 | -0.090 | 1 | -0.258 | 1 | -0.138 | 1 |

**Table S6**: Post-hoc test results for total mineral N recovered ($NH_4^+$, $NO_3^-$ and $NO_2^-$) differences between treatments on each day for (A) the AS and (B) the CS. Pairwise comparisons of estimated marginal means (EMMs) from a gamma-log generalized linear model (glm) were done for both soils. Significant differences relative to the control as well as between treatments are depicted in bold.

A

| Comparison | Day 0 | | Day 2 | | Day 3 | | Day 8 | | Day 15 | |
|---|---|---|---|---|---|---|---|---|---|---|
| | Est. | p adj | Est. | p adj | Est. | p adj | Est. | p adj | Est. | p adj |
| DMPP-Control | 0.142 | 1 | -0.039 | 1 | 0.163 | 1 | 0.122 | 1 | 0.124 | 1 |
| Limonene-Control | 0.047 | 1 | -0.072 | 1 | 0.137 | 1 | **0.597** | **< 0.001** | **0.761** | **< 0.001** |
| MBOA-Control | 0.084 | 1 | -0.067 | 1 | 0.094 | 1 | 0.095 | 1 | -0.046 | 1 |
| MHPA-Control | 0.139 | 1 | 0.012 | 1 | 0.132 | 1 | 0.113 | 1 | 0.076 | 1 |
| Limonene-DMPP | -0.095 | 1 | -0.034 | 1 | -0.026 | 1 | **0.475** | **< 0.001** | **0.638** | **< 0.001** |
| MBOA-DMPP | -0.058 | 1 | -0.029 | 1 | -0.068 | 1 | -0.027 | 1 | -0.170 | 1 |
| MHPA-DMPP | -0.003 | 1 | 0.051 | 1 | -0.031 | 1 | -0.009 | 1 | -0.047 | 1 |
| MBOA-Limonene | 0.037 | 1 | 0.005 | 1 | -0.043 | 1 | **-0.502** | **< 0.001** | **-0.807** | **< 0.001** |
| MHPA-Limonene | 0.092 | 1 | 0.084 | 1 | -0.005 | 1 | **-0.484** | **< 0.001** | **-0.685** | **< 0.001** |
| MHPA-MBOA | 0.055 | 1 | 0.079 | 1 | 0.037 | 1 | 0.018 | 1 | 0.123 | 1 |

B

| Comparison | Day 0 | | Day 3 | | Day 6 | | Day 15 | | Day 21 | |
|---|---|---|---|---|---|---|---|---|---|---|
| | Est. | p adj | Est. | p adj | Est. | p adj | Est. | p adj | Est. | p adj |
| DMPP-Control | 0.045 | 1 | 0.028 | 1 | 0.009 | 1 | **0.260** | **< 0.001** | **0.256** | **0.001** |
| Limonene-Control | 0.089 | 1 | 0.122 | 1 | **0.297** | **< 0.001** | **0.301** | **< 0.001** | **0.332** | **< 0.001** |
| MBOA-Control | -0.009 | 1 | 0.024 | 1 | -0.008 | 1 | 0.168 | 0.478 | 0.020 | 1 |
| MHPA-Control | 0.053 | 1 | 0.069 | 1 | 0.022 | 1 | 0.134 | 1 | 0.039 | 1 |
| Limonene-DMPP | 0.044 | 1 | 0.093 | 1 | **0.288** | **< 0.001** | 0.040 | 1 | 0.076 | 1 |
| MBOA-DMPP | -0.054 | 1 | -0.005 | 1 | -0.016 | 1 | -0.092 | 1 | **-0.236** | **0.005** |
| MHPA-DMPP | 0.008 | 1 | 0.040 | 1 | 0.013 | 1 | -0.126 | 1 | **-0.216** | **0.021** |
| MBOA-Limonene | -0.098 | 1 | -0.098 | 1 | **-0.304** | **< 0.001** | -0.133 | 1 | **-0.312** | **< 0.001** |
| MHPA-Limonene | -0.036 | 1 | -0.053 | 1 | **-0.274** | **< 0.001** | -0.166 | 0.533 | **-0.292** | **< 0.001** |
| MHPA-MBOA | 0.062 | 1 | 0.045 | 1 | 0.030 | 1 | -0.034 | 1 | 0.019 | 1 |

**Table S7**: Post-hoc test results for differences in AOA *amoA* transcript abundances between treatments on each day for (A) the AS and (B) the CS. For the AS, Tukey HSD comparisons were done after a two-way ANOVA, while pairwise comparisons of estimated marginal means (EMMs) from a gamma-log generalized linear model (glm) were used for the CS. Significant differences relative to the control as well as between treatments are depicted in bold.

A

| Comparison | Day 0 | | Day 3 | | Day 8 | | Day 15 | |
|---|---|---|---|---|---|---|---|---|
| | diff | p adj | diff | p adj | diff | p adj | diff | p adj |
| DMPP-Control | 0.171 | 0.999 | 0.015 | 1 | **0.860** | **< 0.001** | 0.122 | 0.999 |
| Limonene-Control | -0.014 | 1 | **-0.992** | **< 0.001** | **-1.168** | **< 0.001** | **-2.430** | **< 0.001** |
| MBOA-Control | 0.421 | 0.614 | **-0.819** | **0.002** | -0.577 | 0.118 | **-1.417** | **< 0.001** |
| MHPA-Control | -0.538 | 0.197 | -0.356 | 0.853 | 0.174 | 0.999 | -0.269 | 0.987 |
| Limonene-DMPP | -0.185 | 0.999 | **-1.01** | **< 0.001** | **-2.028** | **< 0.001** | **-2.552** | **< 0.001** |
| MBOA-DMPP | 0.250 | 0.994 | **-0.834** | **0.001** | **-1.437** | **< 0.001** | **-1.540** | **< 0.001** |
| MHPA-DMPP | -0.709 | 0.014 | -0.371 | 0.808 | **-0.686** | **0.021** | -0.392 | 0.733 |
| MBOA-Limonene | 0.435 | 0.557 | 0.173 | 0.999 | 0.591 | 0.096 | **1.012** | **< 0.001** |
| MHPA-Limonene | -0.524 | 0.233 | 0.636 | 0.050 | **1.342** | **< 0.001** | **2.160** | **< 0.001** |
| MHPA-MBOA | **-0.960** | **< 0.001** | 0.463 | 0.442 | **0.751** | **0.006** | **1.150** | **< 0.001** |

B

| Comparison | Day 0 | | Day 6 | | Day 15 | | Day 21 | |
|---|---|---|---|---|---|---|---|---|
| | Est. | p adj | Est. | p adj | Est. | p adj | Est. | p adj |
| Control-DMPP | 0.002 | 1 | -0.028 | 1 | **-0.256** | **< 0.001** | **-0.261** | **< 0.001** |
| Control-Limonene | 0.042 | 1 | 0.039 | 1 | -0.100 | 1 | -0.025 | 1 |
| Control-MBOA | 0.046 | 1 | 0.011 | 1 | -0.125 | 0.213 | -0.039 | 1 |
| Control-MHPA | 0.027 | 1 | -0.006 | 1 | -0.123 | 0.252 | -0.029 | 1 |
| DMPP-Limonene | 0.040 | 1 | 0.067 | 1 | **0.156** | **0.014** | **0.235** | **< 0.001** |
| DMPP-MBOA | 0.044 | 1 | 0.039 | 1 | 0.131 | 0.135 | **0.222** | **< 0.001** |
| DMPP-MHPA | 0.025 | 1 | 0.022 | 1 | 0.133 | 0.113 | **0.232** | **< 0.001** |
| Limonene-MBOA | 0.004 | 1 | -0.028 | 1 | -0.025 | 1 | -0.013 | 1 |
| Limonene-MHPA | -0.015 | 1 | -0.045 | 1 | -0.023 | 1 | -0.003 | 1 |
| MBOA-MHPA | -0.019 | 1 | -0.017 | 1 | 0.002 | 1 | 0.010 | 1 |

**Table S8**: Post-hoc test results for differences in AOB *amoA* transcript abundances between treatments on each day for (A) the AS and (B) the CS. Pairwise comparisons of estimated marginal means (EMMs) after a gamma generalized linear model (glm) and a gamma log glm were used for the AS and CS, respectively. Significant differences relative to the control as well as between treatments are depicted in bold.

A

| **Comparison** | **Day 0** | | **Day 3** | | **Day 8** | | **Day 15** | |
|---|---|---|---|---|---|---|---|---|
| | **Est.** | **p adj** | **Est.** | **p adj** | **Est.** | **p adj** | **Est.** | **p adj** |
| Control-DMPP | -0.330 | 1 | 1.701 | 0.107 | **4.306** | **< 0.001** | **3.106** | **< 0.001** |
| Control-Limonene | **2.484** | **< 0.001** | **2.845** | **< 0.001** | 0.734 | 1 | -0.913 | 1 |
| Control-MBOA | 0.401 | 1 | 1.496 | 0.446 | 0.794 | 1 | **-2.327** | **< 0.001** |
| Control-MHPA | 0.626 | 1 | 0.533 | 1 | -0.216 | 1 | -1.659 | 0.052 |
| DMPP-Limonene | **2.814** | **< 0.001** | 1.144 | 1 | **-3.572** | **< 0.001** | **-4.019** | **< 0.001** |
| DMPP-MBOA | 0.732 | 1 | -0.204 | 1 | **-3.513** | **< 0.001** | **-5.432** | **< 0.001** |
| DMPP-MHPA | 0.956 | 1 | -1.168 | 1 | **-4.523** | **< 0.001** | **-4.764** | **< 0.001** |
| Limonene-MBOA | **-2.082** | **< 0.001** | -1.348 | 0.334 | 0.060 | 1 | -1.413 | 0.626 |
| Limonene-MHPA | **-1.858** | **< 0.001** | **-2.312** | **< 0.001** | -0.950 | 1 | -0.745 | 1 |
| MBOA-MHPA | 0.225 | 1 | -0.963 | 1 | -1.010 | 1 | 0.668 | 1 |

B

| **Comparison** | **Day 0** | | **Day 6** | | **Day 15** | | **Day 21** | |
|---|---|---|---|---|---|---|---|---|
| | **Est.** | **p adj** | **Est.** | **p adj** | **Est.** | **p adj** | **Est.** | **p adj** |
| Control-DMPP | -0.011 | 1 | **0.263** | **< 0.001** | **0.308** | **< 0.001** | **0.479** | **< 0.001** |
| Control-Limonene | 0.076 | 1 | -0.029 | 1 | -0.049 | 1 | 0.126 | 0.598 |
| Control-MBOA | 0.067 | 1 | 0.002 | 1 | 0.025 | 1 | 0.056 | 1 |
| Control-MHPA | 0.051 | 1 | -0.012 | 1 | -0.054 | 1 | 0.084 | 1 |
| DMPP-Limonene | 0.087 | 1 | **-0.292** | **< 0.001** | **-0.356** | **< 0.001** | **-0.353** | **< 0.001** |
| DMPP-MBOA | 0.078 | 1 | **-0.261** | **< 0.001** | **-0.283** | **< 0.001** | **-0.423** | **< 0.001** |
| DMPP-MHPA | 0.062 | 1 | **-0.276** | **< 0.001** | **-0.362** | **< 0.001** | **-0.395** | **< 0.001** |
| Limonene-MBOA | -0.008 | 1 | 0.031 | 1 | 0.074 | 1 | -0.069 | 1 |
| Limonene-MHPA | -0.024 | 1 | 0.017 | 1 | -0.004 | 1 | -0.042 | 1 |
| MBOA-MHPA | -0.016 | 1 | -0.015 | 1 | -0.078 | 1 | 0.028 | 1 |

**Table S9**: Post-hoc tests results of differences in AOA *amoA* transcript-to gene ratios between treatments on each day for (A) the AS and (B) the CS. For the AS, Tukey HSD comparisons were done after a two-way ANOVA, while pairwise comparisons of estimated marginal means (EMMs) after a gamma-log generalized linear model (glm) were used for the CS. Significant differences relative to the control as well as between treatments are depicted in bold.

A

| Comparison | Day 0 | | Day 3 | | Day 8 | | Day 15 | |
|---|---|---|---|---|---|---|---|---|
| | diff | p adj | diff | p adj | diff | p adj | diff | p adj |
| DMPP-Control | -0.163 | 1 | -0.027 | 1 | **0.952** | **0.002** | 0.196 | 1 |
| Limonene-Control | -0.662 | 0.140 | **-0.868** | **0.008** | **-1.106** | **< 0.001** | **-1.860** | **< 0.001** |
| MBOA-Control | -0.336 | 0.976 | -0.519 | 0.528 | -0.533 | 0.477 | **-0.932** | **0.003** |
| MHPA-Control | -0.665 | 0.136 | -0.226 | 1 | 0.157 | 1 | -0.106 | 1 |
| Limonene-DMPP | -0.500 | 0.596 | **-0.841** | **0.012** | **-2.057** | **< 0.001** | **-2.056** | **< 0.001** |
| MBOA-DMPP | -0.174 | 1 | -0.491 | 0.626 | **-1.485** | **< 0.001** | **-1.128** | **< 0.001** |
| MHPA-DMPP | -0.502 | 0.587 | -0.199 | 1 | **-0.794** | **0.024** | -0.302 | 0.992 |
| MBOA-Limonene | 0.326 | 0.983 | 0.350 | 0.965 | 0.573 | 0.347 | **0.928** | **0.003** |
| MHPA-Limonene | -0.003 | 1 | 0.642 | 0.175 | **1.263** | **< 0.001** | **1.754** | **< 0.001** |
| MHPA-MBOA | -0.329 | 0.981 | 0.292 | 0.995 | 0.690 | 0.100 | **0.826** | **0.015** |

B

| Comparison | Day 0 | | Day 6 | | Day 15 | | Day 21 | |
|---|---|---|---|---|---|---|---|---|
| | Est. | p adj | Est. | p adj | Est. | p adj | Est. | p adj |
| Control-DMPP | 0.030 | 1 | -0.080 | 1 | **-1.704** | **< 0.001** | **-2.534** | **< 0.001** |
| Control-Limonene | 0.209 | 1 | 0.682 | 1 | 0.378 | 1 | -0.338 | 1 |
| Control-MBOA | 0.053 | 1 | 0.438 | 1 | 0.035 | 1 | -0.148 | 1 |
| Control-MHPA | 0.142 | 1 | 0.179 | 1 | -0.360 | 1 | -0.032 | 1 |
| DMPP-Limonene | 0.179 | 1 | 0.762 | 0.992 | **2.082** | **< 0.001** | **2.196** | **< 0.001** |
| DMPP-MBOA | 0.023 | 1 | 0.518 | 1 | **1.739** | **< 0.001** | **2.386** | **< 0.001** |
| DMPP-MHPA | 0.111 | 1 | 0.259 | 1 | **1.343** | **< 0.001** | **2.501** | **< 0.001** |
| Limonene-MBOA | -0.155 | 1 | -0.244 | 1 | -0.343 | 1 | 0.190 | 1 |
| Limonene-MHPA | -0.067 | 1 | -0.503 | 1 | -0.739 | 1 | 0.305 | 1 |
| MBOA-MHPA | 0.088 | 1 | -0.259 | 1 | -0.396 | 1 | 0.115 | 1 |

**Table S10**: Post-hoc tests results of differences in AOB *amoA* transcript-to gene ratios between treatments on each day for (A) the AS and (B) the CS. Pairwise comparisons of estimated marginal means (EMMs) after a gamma-log generalized linear model (glm) were used for both soils. Significant differences relative to the control as well as between treatments are depicted in bold.

A

| Comparison | Day 0 | | Day 3 | | Day 8 | | Day 15 | |
|---|---|---|---|---|---|---|---|---|
| | Est. | p adj | Est. | p adj | Est. | p adj | Est. | p adj |
| Control-DMPP | -0.330 | 1 | **0.869** | **0.038** | **2.822** | **0.019** | 1.713 | 1 |
| Control-Limonene | 2.894 | 0.086 | **1.844** | **< 0.001** | -0.467 | 1 | **-2.448** | **< 0.001** |
| Control-MBOA | 0.799 | 1 | 0.466 | 1 | -0.041 | 1 | **-2.291** | **< 0.001** |
| Control-MHPA | 0.662 | 1 | 0.134 | 1 | -0.564 | 1 | **-2.177** | **0.002** |
| DMPP-Limonene | **3.224** | **0.016** | 0.975 | 1 | **-3.289** | **< 0.001** | **-4.162** | **0.031** |
| DMPP-MBOA | 1.129 | 0.124 | -0.402 | 1 | **-2.863** | **0.014** | -4.005 | 0.055 |
| DMPP-MHPA | 0.993 | 0.317 | -0.735 | 0.388 | **-3.386** | **< 0.001** | -3.890 | 0.083 |
| Limonene-MBOA | -2.095 | 1 | **-1.377** | **0.014** | 0.426 | 1 | 0.157 | 1 |
| Limonene-MHPA | -2.231 | 1 | **-1.710** | **< 0.001** | -0.097 | 1 | 0.271 | 1 |
| MBOA-MHPA | -0.136 | 1 | -0.333 | 1 | -0.523 | 1 | 0.114 | 1 |

B

| Comparison | Day 0 | | Day 6 | | Day 15 | | Day 21 | |
|---|---|---|---|---|---|---|---|---|
| | Est. | p adj | Est. | p adj | Est. | p adj | Est. | p adj |
| Control-DMPP | -0.154 | 1 | **1.166** | **0.008** | **1.954** | **< 0.001** | **1.940** | **< 0.001** |
| Control-Limonene | 0.141 | 1 | -0.387 | 1 | 0.531 | 1 | -0.039 | 1 |
| Control-MBOA | 0.096 | 1 | 0.054 | 1 | 0.640 | 1 | -0.066 | 1 |
| Control-MHPA | 0.047 | 1 | -0.063 | 1 | 0.316 | 1 | 0.687 | 1 |
| DMPP-Limonene | 0.296 | 1 | **-1.553** | **< 0.001** | **-1.423** | **< 0.001** | **-1.980** | **< 0.001** |
| DMPP-MBOA | 0.250 | 1 | **-1.112** | **0.017** | **-1.314** | **0.001** | **-2.007** | **< 0.001** |
| DMPP-MHPA | 0.201 | 1 | **-1.229** | **0.004** | **-1.637** | **< 0.001** | **-1.253** | **0.002** |
| Limonene-MBOA | -0.046 | 1 | 0.441 | 1 | 0.108 | 1 | -0.027 | 1 |
| Limonene-MHPA | -0.095 | 1 | 0.323 | 1 | -0.214 | 1 | 0.726 | 1 |
| MBOA-MHPA | -0.049 | 1 | -0.117 | 1 | -0.323 | 1 | 0.754 | 1 |

**Table S11**: Differences in NI recovery at the start of the incubation period. Data are presented as means ± SE (n = 4).

| | AS | | | CS | | |
|---|---|---|---|---|---|---|
| | Expected concentra-tion (µg $g^{-1}$ soil dw) | Recovered NI amount (µg $g^{-1}$ soil dw) | Recovered NI amount (%) | Expected concentration (µg $g^{-1}$ soil dw) | Recovered NI amount (µg $g^{-1}$ soil dw) | Recovered NI amount (%) |
| DMPP | 1.3 | 0.5 ± 0.02 | 38.0 ± 1.7 | 1.3 | 0.46 ± 0.04 | 35.3 ± 2.9 |
| Limonene | 1800 | 288.7 ± 25.7 | 16.0 ± 1.4 | 786 | 27.2 ± 2.66 | 3.4 ± 0.3 |
| MBOA | 290 | 161.6 ± 12.8 | 55.7 ± 4.4 | 111 | 50.03 ± 4.66 | 45.1 ± 4.2 |
| MHPA | 387 | 211.4 ± 36.2 | 54.6 ± 9.4 | 394 | 119.1 ± 3.8 | 30.2 ± 6.1 |

**Supplementary methods**

**LC-HRMS measurements of BNIs in acetonitrile extracts and DMPP in HPLC eluent**

For the quantification of MHPA, MBOA and DMPP, an IQ-X Orbitrap (Thermo Fisher Scientific, San Jose, CA, USA) equipped with a heated electro-spray ionization source (H-ESI) coupled to a Vanquish Horizon UHPLC system (Thermo Fisher Scientific, San Jose, CA, USA) was used. A reversed phase HPLC Method utilizing an Acquity UPLC® HSS T3 150 × 2.1 mm i.d., 1.8 µm particle size column (Waters, Milford, MA, USA) coupled with a VanGuard FIT Cartridge was used for chromatographic separation.

Operating conditions were as follows: The column temperature and pre-column heater were set to 35 °C and a sample injection volume of 5 µL was defined. Liquid phase was applied at a constant flow rate of 275 $\mu L \cdot min^{-1}$ and consisted of ELGA $H_2O$ with 0.1% formic acid ($v/v^{-1}$) (eluent A) and MeOH with 0.1% formic acid ($v/v^{-1}$) (eluent B). A gradient elution was applied starting with 10 % B for 0.6 min with a subsequent 12.4 min linear increase to 100% B followed by 100% B for 2.5 min and finally a re-equilibration at 10% B for 3.5 min (total run time 19 min per injection).

Orbitrap measurements of MBOA and MHPA were done in full-scan data acquisition mode using the following scan parameters: scan range: *m/z* 140–550, normalized AGC target 50%, maximum injection time 502 ms, and resolving power R = 120,000 at m/z 200. The H-ESI probe was operated in fast polarity switching mode using the following parameter settings: sheath gas flow rate 45 arb. units; Aux gas flow rate 10 arb. units using a vaporizer temperature of 300°C and an ion transfer tube temperature of 270 °C. The Spray voltage was set to 3.25 kV (positive mode) or 3.05 kV (negative mode). For the DMPP the orbitrap settings consisted of full-scan data acquisition with scan range *m/z* 80-1000, tSIM data acquisition with scan range *m/z* 94-100 from retention time 2.5 min to 5.5 min., normalized AGC target 50%, maximum injection time 502 ms, and resolving power R = 240,000 at m/z 200. The H-ESI probe was operated in positive only mode using the following parameter settings: sheath gas flow rate 45 arb. units; Aux gas flow rate 10 arb. Units; vaporizer temperature of 290°C and ion transfer tube temperature 270 °C. The Spray voltage was set to 2.85 kV.

**GC-MS measurements of n-hexane extracts (limonene)**

The concentration of limonene in the n-hexane extracts was quantified following the method established by Huang et al. (2022). Briefly, samples were separated on an HP-5MS capillary column (60 m length, 0.25 mm inner diameter, 0.25 µm film thickness, 5% phenyl-methyl polysiloxane) using high-purity helium as a carrier gas. The initial column temperature was set at 90 °C for 5 min, then it was raised to 230 °C at a rate of 10 °C $min^{-1}$ and maintained at 270 °C for 1 min. The injection volume was 1 μL, and the column flow rate was set to 1.5 mL $min^{-1}$. The ionization energy was 70 eV, and the solvent delay was set to 6.5min.

**LC-HRMS and GC-MS Data Processing**

For quantifying the analytes (NIs) in the extracts, external calibration with eight concentration levels of each substance dissolved in the respective extraction solvent was included. The peak detection parameters for the acetonitrile extracts were as follows: ICIS peak picking algorithm, nearest rt, smoothing points 3, baseline window 40, area noise factor 4 and peak noise factor 9. Data evaluation was done with the Thermo Fisher Scientific Software Xcalibur QuanBrowser (Version 4.7.69.37)

**NI recovery calculations**

Initially, the μg of NI per gram of soil (μg NI $g^{-1}$ dw soil) recovered at each time point were calculated by multiplying the NI concentration determined with LC-HRMS by the extract volume (V) (0.004L for MHPA and MBOA, 0.002L for DMPP) and dividing it by the soil dry weight:

$$\frac{\mu g\ NI}{g\ dw\ soil} = \frac{\left(\frac{\mu g\ NI}{L}\right) \times V}{g\ dw\ soil}$$

For the percentage of NI recovered, the average μg NI $g^{-1}$ dry weight soil from the four soil replicates of each treatment on day 0 was taken as 100%, and the amount of NI recovered on subsequent days was calculated relative to this initial value.
